# Cross species multi-omics reveals cell wall sequestration and elevated global transcription as mechanisms of boron tolerance in plants

**DOI:** 10.1101/2020.10.01.321760

**Authors:** Guannan Wang, Sandra Feuer DiTusa, Dong-Ha Oh, Achim D. Herrmann, David G. Mendoza-Cozatl, Malcolm A. O’Neill, Aaron P. Smith, Maheshi Dassanayake

**Affiliations:** Department of Biological Sciences, Louisiana State University, Baton Rouge, LA, USA; Department of Geology & Geophysics and Coastal Studies Institute, Louisiana State University, Baton Rouge, LA, USA; Division of Plant Sciences, Interdisciplinary Plant Group, Christopher S. Bond Life Sciences Center, University of Missouri, Columbia, MO, USA; Complex Carbohydrate Research Center, The University of Georgia, Athens, GA, USA

**Keywords:** *Schrenkiella parvula*, extremophyte, boron toxicity, boron tolerance, boron transporters, cell wall, RNA metabolism, stress-preparedness, excess boron stress, transcriptome, ionome, metabolome

## Abstract

Boron toxicity is a worldwide problem for crop production, yet we have only a limited understanding of the genetic responses and adaptive mechanisms to this environmental stress in plants. Here we identified responses to excess boron in boron stress-sensitive *Arabidopsis thaliana* and its boron stress-tolerant extremophyte relative *Schrenkiella parvula* using comparative genomics, transcriptomics, metabolomics, and ionomics*. S. parvula* maintains a lower level of total boron and free boric acid in its roots and shoots and sustains growth for longer durations than *A. thaliana* when grown with excess boron. *S. parvula* likely excludes boron more efficiently than *A. thaliana*, which we propose is partly driven by BOR5, a boron transporter that we functionally characterized in the current study. Both species allocate significant transcriptomic and metabolomic resources to enable their cell walls to serve as a partial sink for excess boron, particularly discernable in *A. thaliana* shoots. We provide evidence that the *S. parvula* transcriptome is pre-adapted to boron toxicity, exhibiting substantial overlap with the boron-stressed transcriptome of *A. thaliana*. Our transcriptomic and metabolomics data also suggest that RNA metabolism is a primary target of boron toxicity. Cytoplasmic boric acid likely forms complexes with ribose and ribose-containing compounds critical to RNA and other primary metabolic functions. A model depicting some of the cellular responses that enable a plant to grow in the presence of normally toxic levels of boron is presented.

## Introduction

In 1899 the eminent botanist Edwin Copeland stated that “boron produced monstrosity” to describe plant damage due to excess boron (Copeland and Kahlenberg, 1899). Boron functions as an essential micronutrient in plants at a narrow concentration range (0.5 - 1 ppm, equivalent to 46.2 - 92.5 μM in hydroponic media), causing severe growth defects in many plants, including most crops, at only slightly higher concentrations (Brenchley, 1914; Haas, 1929; Warington, 1937; Eaton, 1940; Goldberg, 1997; Reid, 2007, 2013; Julkowska, 2018; Landi et al., 2019). Early surveys of boron toxicity effects led plants to be classified as sensitive, semi-tolerant, or tolerant to boron (Eaton, 1935). Subsequently, the Food and Agriculture Organization of the United Nations recommended a soil boron level of less than 1.4 mM for even the most tolerant crops to minimize losses in productivity (Eaton, 1944; Ayers and Westcot, 1985; Grieve et al., 2011). The negative impact of boron toxicity on US agriculture was recognized early on (Cook and Wilson, 1918; Eaton, 1935). It is also known to reduce crop yields on all continents where agricultural regions are affected by naturally high amounts of boron in soils or in irrigation water, particularly when the water is obtained from sources near active geothermal areas (Nable et al., 1997; Camacho-cristóbal et al., 2008; Reid and Fitzpatrick, 2009b). Moreover, most soils containing toxic levels of boron occur in semi-arid environments where drought and high salinity compound the stresses on the crops (Reid, 2010).

Excess boron inhibits plant growth by decreasing chlorophyll content, stomatal conductance, photosynthesis, and leads to premature death of shoots and roots (Lovatt and Bates, 1984; Reid et al., 2004; Miwa et al., 2007). Nevertheless, the molecular targets of excess boron and the cellular and molecular processes interrupted by boron stress are poorly understood. Similarly, we have little understanding of the genetic mechanisms underlying boron toxicity responses or the adaptive mechanisms plants use to counter excess boron (Reid et al., 2004; Ruiz et al., 2003; Princi et al., 2016).

In this study, we used the boron-tolerant extremophyte *Schrenkiella parvula* (formerly *Thellungiella parvula* and *Eutrema parvulum*, family Brassicaceae) (Dassanayake et al., 2011; Zhu, 2015; Kazachkova et al., 2018) and its close relative *A. thaliana*, a boron-sensitive model, to identify cellular processes interrupted by excess boron and to determine the transcriptional and metabolic processes that support growth during boron toxicity. *S. parvula* is adapted to high levels of boron naturally present in its native habitats in the Central Anatolian plateau of Turkey (Helvaci et al., 2004). The ecotype (Lake Tuz) used in our study was collected from the Lake Tuz region of Turkey and experiences an average concentration of boron (2.2 mM) that is highly toxic to most plants (Nilhan et al., 2008). It can survive soil boron levels as high as 5.8 mM boron in the wild (Nilhan et al., 2008) and 10 mM boron given for two weeks in controlled environments (Oh et al., 2014). However, the mechanisms that allow *S. parvula* to grow in the presence of boron concentrations that are toxic to *A. thaliana* are not known.

Here, we used comparative genomics, transcriptomics, ionomics, and metabolomics to study boron toxicity responses and tolerance in *A. thaliana* and *S. parvula*. Our data suggest that excess boron disturbs cell wall metabolism and RNA metabolism-related processes, particularly translation. The cell walls of *A. thaliana* and *S. parvula* serve as a sink to partially sequester excess boron under high boron conditions. *S. parvula* accumulated less boron than *A. thaliana* under boron toxicity, likely through an efficient efflux system. We propose that *S. parvula* has a pre-adapted transcriptome to facilitate rapid metabolic changes when exposed to excess boron and that such a pre-adaptation distinguishes boron stress-adapted and -sensitive plants. We provide a model depicting critical cellular processes that are affected by excess boron and the molecular mechanisms boron stress-tolerant plants use to minimize the growth inhibitory effects of this element.

## Results

### *S. parvula* accumulates less boron while sustaining growth for longer durations compared to *A. thaliana*

*S. parvula* was unaffected by treatments of 5, 10, or 15 mM boric acid whether grown hydroponically for four weeks (Figure 1A) or on plates for one week (Figure 1B). In contrast, *A. thaliana* showed clear growth inhibition, wilting, and chlorosis of leaves 5 or 7 days after the boric acid treatments (Figure 1A and B). The control growth media included ~100 μM boron to provide a boron-sufficient growth medium, and the treatments added excess boron to this growth-sufficient level. We observed a substantial reduction in the fresh and dry weights of *A. thaliana* shoots and roots in response to excess boron, whereas *S. parvula* biomass was unaffected by the treatments (Figure 1C). Similarly, total chlorophyll content decreased in *A. thaliana* but remained unchanged in *S. parvula* at 3 days after the treatments (Figure 1D). In *A. thaliana*, we observed dose-dependent inhibitory effects of excess boron on root growth, whereas *S. parvula* root growth was not affected (Figure 1E). In *A. thaliana*, the 15 mM boron treatment led to a significant reduction of lateral root density (Figure 1F), while average lateral root length was decreased in all treatments (Figure 1G). In contrast, neither lateral root density nor average lateral root length of *S. parvula* was affected by excess boron (Figure 1F and G).

**Figure 1.**
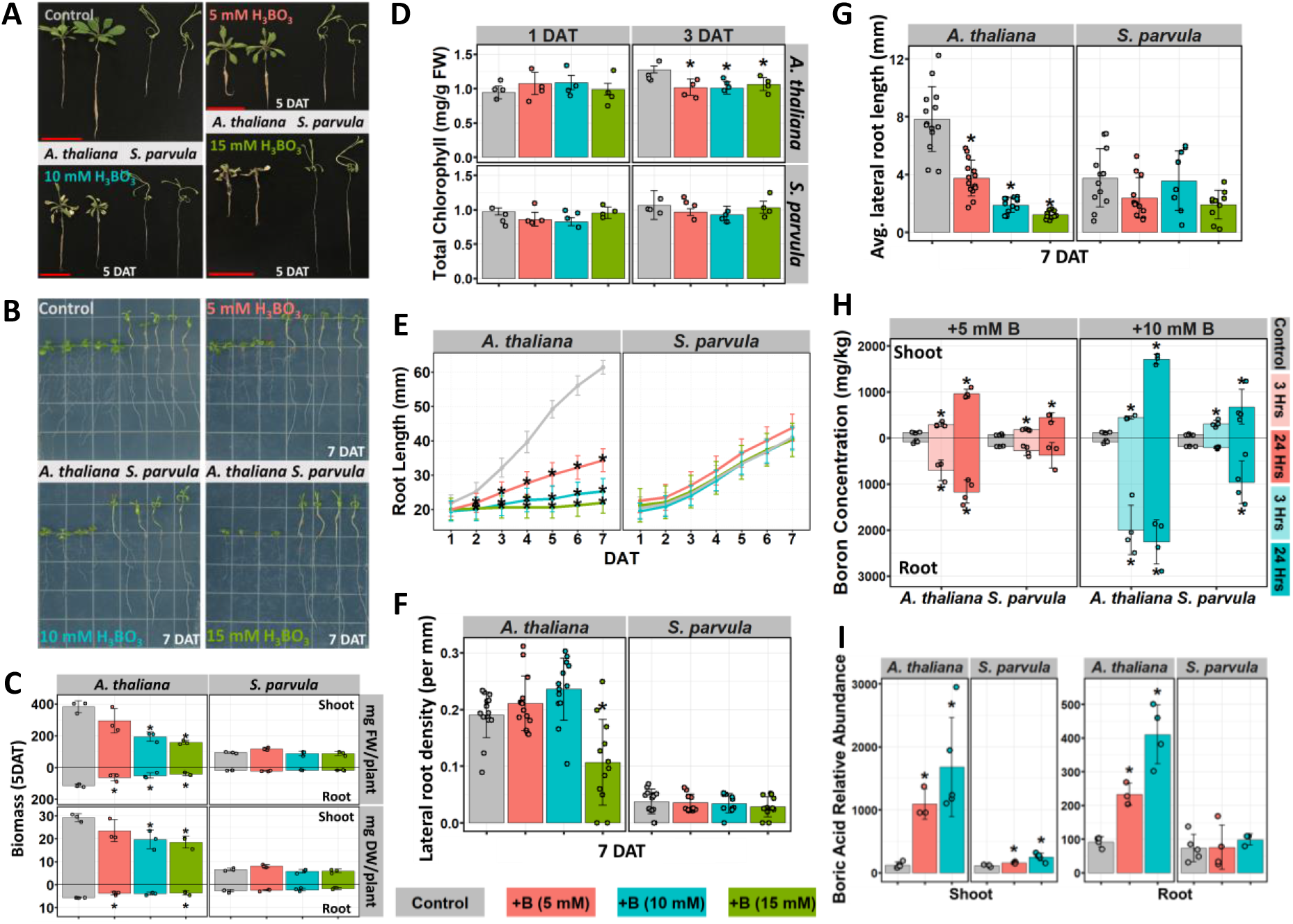
*A. thaliana* and *S. parvula* respond to boric acid treatments differently. (A) Hydroponically grown 33-day-old *A. thaliana* and *S. parvula* with different concentrations of boric acid. Boric acid treatments started at 4-week-old plants. Scale bars = 5 cm. (B) Growth phenotype of 2-week-old *A. thaliana* and *S. parvula* on plates with boric acid. Plants were germinated and grown on 1/4 MS medium, transferred to 1/4 MS medium supplemented with boric acid one week after the germination. (C) Dry biomass and (D) total chlorophyll content of hydroponically grown *A. thaliana* and *S. parvula*. (E) Root growth, (F) lateral root density, and (G) average lateral root length of plate-grown *A. thaliana* and *S. parvula* seedlings. (H) Boron and (I) free boric acid accumulation in shoots and roots in *A. thaliana* and *S. parvula*. In panels C-I, all values are mean ± SD (n=3~5, except for E where n=14~15). Asterisks represent significant differences (p<0.05) compared to control determined by either one-way ANOVA followed by Tukey’s post-hoc tests (C-H) or Student’s t-test (I).

We quantified the concentration of boron (on a dry weight basis) in shoots and roots of plants exposed to excess boron using inductively coupled plasma mass spectrometry (ICP-MS) to determine if the distinct boron stress-responses between *S. parvula* and *A. thaliana* reflected differences in their boron accumulation (Figure 1H). The initial boron levels in roots and shoots of control plants were similar for both species. Differences in boron accumulation only differed in response to the excess boron treatments. The levels of boron increased significantly in the roots and shoots of both species over time and with higher concentrations of boron. However, the level of boron that accumulated in *S. parvula* was lower than in *A. thaliana*. This was most apparent in roots, 24 hours after the 5 mM boric acid treatment, in which boron levels increased 14-fold in *A. thaliana* but were unchanged in *S. parvula* (Figure 1H). The greater capacity of *S. parvula* to maintain lower boron levels relative to *A. thaliana* may have contributed to the continued root growth in *S. parvula* under conditions that decreased *A. thaliana* root growth (Figure 1E). However, *S. parvula* roots could not limit boron accumulation comparable to control plants when grown on 10 mM boric acid. Nevertheless, under this treatment, the relative accumulation of boron in roots was still much lower in *S. parvula* than in *A. thaliana* (Figure 1H). In contrast to roots, *S. parvula* shoots significantly accumulated boron even within 3 hours of the 5 mM boric acid treatment, although the amounts were lower than in *A. thaliana* shoots under all comparable treatments (Figure 1H). This indicated that the overall response to excess boron is different between roots and shoots.

We next determined if the physiological responses (Figure 1 A-G) to excess boron coincided with a disruption in the ionic balance of other elements. Reduction of nutrient uptake or concurrent over-accumulation of other elements may cause toxicity symptoms not directly attributed to boron stress. To this end, we quantified 20 other elements known for their presence in plants (Supplemental Figure 1). Neither species showed dramatic changes to their ionomic profiles during boron treatments except for the expected increase in boron content, suggesting that the observed physiological responses in both species were largely caused by the cellular disturbances due to excess boron accumulation.

### High external boron results in a substantial increase in free boric acid in *A. thaliana* roots but not in *S. parvula* roots

After being transported primarily as free boric acid into the cytoplasm, boron is either found as free boric acid/borate or bound to organic metabolites within plant cells (Woods, 1996; Hu and Brown, 1997; Brown et al., 2002; Broadley et al., 2012; Camacho-cristóbal et al., 2008). We therefore measured the relative abundance of free boric acid in plants using gas chromatography-mass spectrometry (GC-MS), to assess how *S. parvula* may retain externally supplied boric acid differently from *A. thaliana* (Figure 1I). Notably, we detected substantial increases (up to 14 fold) in free boric acid in treated *A. thaliana* shoots compared to the control, which was comparable to the increase in total boron accumulation (Figure 1H). In contrast, the increase in free boric acid (up to 4.5 fold, Figure 1I) was much lower than the total boron accumulation in *A. thaliana* roots (Figure 1H). This suggested that a major fraction of the increased total boron in *A. thaliana* roots was in the form of B-complexes. *S. parvula* shoots were similar to *A. thaliana* roots since the increase in free boric acid (up to 2.2 fold; Figure 1I) was much lower than the increase of total boron in treated *S. parvula* shoots (Figure 1H). It is equally noteworthy that the levels of free boric acid remained unchanged in *S. parvula* roots under all conditions tested (Figure 1I) despite the substantial increases of total boron in plants grown on 10 mM boric acid (Figure 1H). This led us to hypothesize that *S. parvula* stores much of the excess boron in the form of B-complexes and thereby minimizes the accumulation of free boric acid, particularly in roots, more effectively than *A. thaliana*.

### The transcriptomic response to excess boron is greater in *A. thaliana* than *S. parvula*

We expected that *A. thaliana* and *S. parvula*, which have rapid yet divergent responses to excess boron, would exhibit regulation at the transcriptional level that determined their subsequent responses to this stress. To develop a comparative framework to contrast transcriptional responses to excess boron, plants were transferred to media containing 5 mM boric acid. This concentration was sufficient to induce discernible changes in both species (Figure 1), but did not cause tissue death in the sensitive model even at 5 days post treatment (Figure 1A). We chose a 24-hour duration to allow us to assess transcriptomic responses when neither species showed observable changes to suggest cell death. Since the root stress response was different from that of shoots, we investigated root and shoot transcriptomes separately in our comparative–omics framework.

The largest observed variance (>40%) in the transcriptomes within a species was attributed to the tissue differences (i.e. root versus shoot) as seen in the principal component analysis (PCA) (Figure 2A). However, when we compared the transcriptomes of the same tissues across species, the treated *A. thaliana* transcriptomes were strikingly different from the control, whereas *S. parvula* treated and control transcriptomes were almost indistinguishable (Figure 2B). This suggested that the extent of the transcriptional adjustment to excess boron is much greater in *A. thaliana* than in *S. parvula*. Indeed, we found 9,657 genes in shoots and 6,126 genes in roots differentially expressed in response to excess boron in *A. thaliana* (Supplemental Figure 2). In contrast, the number of boron stress-responsive genes in *S. parvula* was much smaller (535 in shoots and 63 in roots) (Supplemental Figure 2, Supplemental Data Set 1). The magnitude of the differences in the overall transcriptomes reflected the visible physiological responses of these two species to excess boron (Figure 1).

**Figure 2.**
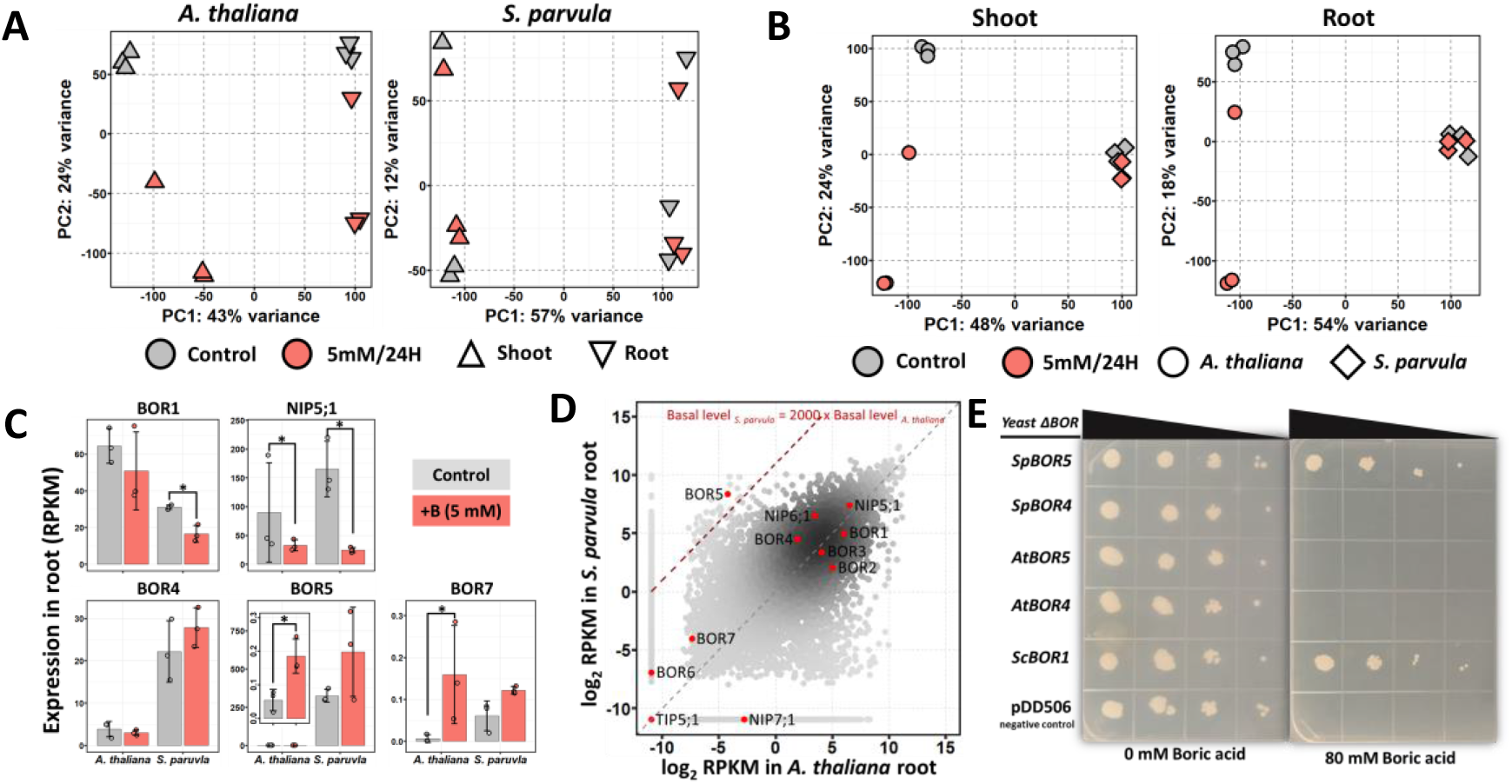
Transcriptional responses of *A. thaliana* and *S. parvula* to excess boron. (A) Principal component analysis (PCA) differentiates the transcriptomes of control and treated samples from shoot and root tissues of *A. thaliana* (left) and *S. parvula* (right). (B) PCA of *A. thaliana* and *S. parvula* transcriptomes within shoot (left) and root (right). (C) Expression levels of BOR1, NIP5;1, BOR4, BOR5, BOR7 in control and 5 mM boric acid treatment. Asterisks represent significant differences in expression compared to control (at FDR-adjusted p<0.05) determined by both DESeq2 and NOISeq. (D) Comparison of the basal expression levels of ortholog pairs in roots between *A. thaliana* and *S. parvula*. Ortholog pairs that encode boron transporters and channels are marked in red. Gray diagonal dashed line marks identical basal level expression between the two species while ortholog pairs above the red dashed line show >2000-times higher basal expression in *S. parvula* than in *A. thaliana*. (E). Growth of yeast *Δbor* mutants transformed with either *ScBOR1*, *AtBOR4*, *AtBOR5*, *SpBOR4*, or *SpBOR5* on medium containing 0 and 80 mM boric acid. Negative control was transformed with the empty vector.

To independently assess the reproducibility of the transcriptomic responses captured by RNAseq, we selected five to six differentially expressed genes per tissue in both species and additional biological replicates to obtain the relative expression of 20 genes using RT-qPCR. We found high concordance in transcript level changes between the RNAseq and RT-qPCR data (Pearson *R*^2^ = 0.71, p = 2.65e-07) (Supplemental Figure 3).

### *A. thaliana* and *S. parvula* exhibit differences in boron transporter gene expression and function

Boron uptake and translocation are known to involve a family of boron transporters (BORs) and membrane intrinsic proteins (MIPs) (Diehn et al., 2019; Yoshinari and Takano, 2017). We compared the expression changes of all known boron transporters and channels to determine the transcriptional responses to excess boron related to boron acquisition and transport (Supplemental Data Set 1, Figure 2C, D). *S. parvula* roots showed down-regulation of both *NIP5;1* and *BOR1*, the two dominant transporters mediating boron uptake and xylem loading during boron-deficient conditions (Takano et al., 2002, 2006). *A. thaliana* roots also showed down-regulation of *NIP5;1* (Figure 2C). This observation supports the idea that both species attempted to reduce boron uptake as well as boron transport to the shoot in response to excess boron.

Plants may alleviate boron toxicity by activating transporters that export boron back to soil or to other compartments away from actively growing tissues, in addition to minimizing boron uptake. BOR4 is the only boron transporter demonstrated to alleviate boron toxicity by moving excess boron from roots back to soil (Miwa et al., 2014, 2007). Our transcriptomic data showed that *BOR4* transcript abundance was not affected by excess boron in either *A. thaliana* or *S. parvula* (Figure 2C). However, basal transcript levels of *SpBOR4* were ~5 fold higher (22.2 RPKM) than those of *AtBOR4* (3.8 RPKM) in roots (Figure 2C, D). We detected two under-studied putative boron transporters (*BOR5* and *BOR7*) that were significantly induced by excess boron. One of these, BOR5 is the closest homolog of BOR4 (Takano et al., 2008; Sun et al., 2012; Oh and Dassanayake, 2019). Whereas *BOR5* was induced (~3 fold) by excess boron in *A. thaliana* roots (Figure 2C), the *S. parvula* ortholog showed a dramatically higher constitutive expression prior to the stress and represents one of the largest basal expression differences in roots (>2000 fold higher in *S. parvula*) observed among all ortholog pairs between the two species (Figure 2D). *BOR7*, encoding another BOR4-like boron transporter (Luo et al., 2019), was also induced in *A. thaliana* roots in response to excess boron, suggesting a putative function to exclude boron under excess boron conditions (Figure 5A). However, *BOR7* was hardly detected at basal levels (lower than 0.1 RPKM) in *A. thaliana* or *S. parvula* roots (Figure 2C). Taken together, our data suggest that, under toxic levels of boron, *A. thaliana* induced the transcript levels for boron transporters implicated in boron exclusion from the roots. On the other hand, in *S. parvula* roots, both *SpBOR4* and *SpBOR5*, although not induced by excess boron, were expressed at a basal level much higher than their orthologs in *A. thaliana*. Therefore, we hypothesized that BOR5 is functionally active in excluding boron in *S. parvula* roots exposed to excess boron.

To further assess the role of SpBOR5 as a key contributor for boron exclusion in *S. parvula*, we individually expressed *BOR4* and *BOR5* from each species in a yeast mutant lacking the native boron transporter *ScBOR1* (Figure 2E). This *Δbor* yeast mutant is sensitive to high concentrations of boric acid because of its inability to export excess boron. *SpBOR5* fully rescued the growth of *Δbor* yeast exposed to a toxic level of boron, while *AtBOR4*, *AtBOR5*, or *SpBOR4* failed to complement the *Δbor* mutant growth defects (Figure 2E). This demonstrates that *SpBOR5* functions similar to *ScBOR1* and seems to have a higher boron efflux capacity compared to *AtBOR4*, *AtBOR5*, and *SpBOR4*. These results, together with the strikingly higher basal expression of *SpBOR5* in *S. parvula* roots, suggest that SpBOR5 likely enabled *S. parvula* to exclude excess boron more efficiently than its stress-sensitive relative *A. thaliana*.

### Transcriptomic responses to boron predict altered cell wall metabolism as a major cellular response to enable cell walls to capture excess boron

We next examined biological processes and functions enriched among the differentially expressed genes (DEGs) in *A. thaliana* roots as it exhibits readily discernible responses when treated with excess boron (Supplemental Data Set 1). From a total of 3,504 boron stress-repressed DEGs, 2,728 could be associated with specific GO functions (Supplemental Data Set 2). Among them, we identified 19 functional clusters using GOMCL (Wang et al., 2020). In short, GOMCL clustered enriched GO terms that had shared genes (>50%) using Markov Clustering and identified non-redundant representative functions within a GO network. The top ten root clusters included over 96% of the GO-annotated DEGs (Figure 3A). This approach revealed that cell wall-related processes account for the largest proportion among boron stress-suppressed DEGs in *A. thaliana* roots. Three of the top ten functional clusters (C2, 7, and 8) were associated with cell wall-related processes, accounting for 1,453 DEGs (Figure 3A, red boxes; Supplemental Data Set 2).

**Figure 3.**
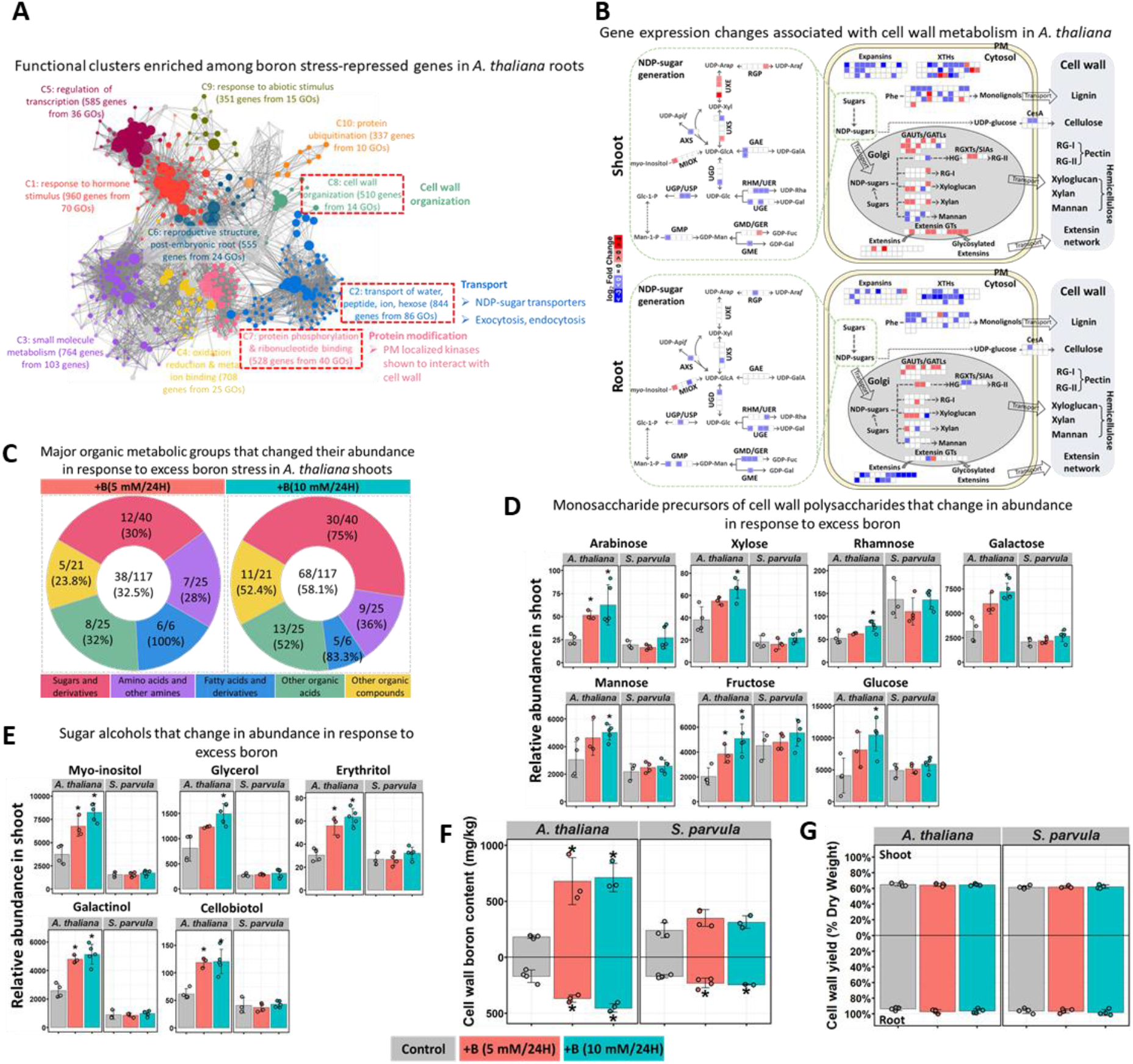
Cell wall metabolism is altered under boron toxicity in *A. thaliana*. (A) Functional clusters enriched among boron stress-repressed genes in *A. thaliana* roots. Clusters associated with cell wall metabolism are marked by red-dashed boxes. Clusters are differently colored and labelled with the representative functional term. Each node represents a GO term; node size represents genes in the test set assigned to that functional term; GO terms sharing more than 50% of genes are connected with edges; and shade of each node represents the *p*-value assigned by the enrichment test (FDR-adjusted p<0.05) with darker shades indicating smaller *p*-values. (B) Changes in gene expression associated with biosynthesis of cell wall components in response to boric acid treatment in *A. thaliana* shoots and roots. Genes are represented by square blocks, grouped by families or pathways, with up- and down-regulation marked by red and blue, respectively. UDP, uridine diphosphate; UDP-Glc, UDP-glucose; UDP-Gal, UDP-galactose; UDP-GlcA, UDP-glucuronic acid; UDP-GalA, UDP-galacturonic acid; UDP-Xyl, UDP-xylose; UDP-Ara, UDP-arabinose; UDP-Araf, UDP-arabinofuranose; UDP-Rha, UDP-rhamnose; UDP-Api, UDP-apiose; GDP-Man, GDP-mannose; GDP-Fuc, GDP-fucose; GDP-Gal, GDP-galactose; GAE, UDP-D-glucuronic acid 4-epimerase; UXS, UDP-D-xylose synthase; UXE, UDP-D-xylose 4-epimerase; RGP, reversibly glycosylated protein; AXS, UDP-D-apiose/UDP-D-xylose synthase (also known as UAXS); MIOX, inositol oxygenase; UGD, UDP-D-glucose dehydrogenase; UGP/USP, UDP-glucose pyrophosphorylase/UDP-sugar pyrophosphorylase; RHM/UER, rhamnose synthase gene/ nucleotide-rhamnose epimerase-reductase; UGE, UDP-D-glucose 4-epimerase; GMP, GDP-D-mannose pyrophosphorylase; GMD/GER, GDP-D-mannose-4,6-dehydratase/GDP-4-keto-6-deoxy-D-mannose-3,5-epimerase-4-reductase; GME, GDP-D-mannose 3,5-epimerase; EXTs, extensins; EXT GTs, extensin glycosyltransferases; EXPs, expansins; XTHs, xyloglucan endotransglucosylase/hydrolases; RGXT, rhamnogalacturonan xylosyltransferase. (C) Major organic metabolic groups that changed their abundance in response to excess boron stress in *A. thaliana* shoots. For each category, the number and proportion of metabolites that changed in abundance compared to the total identified are shown. (D-E) Monosaccharide precursors of cell wall polysaccharides (D) and sugar alcohols (E) that changed in abundance in shoots of *A. thaliana* and *S. parvula* 24 hours after boric acid treatments. The relative abundance is given compared to the internal standard, ribitol. (F) Boron contents in cell walls extracted from shoots and roots of *A. thaliana* and *S. parvula* under different treatments for 24 hours. (G) Cell wall yield of shoots and roots in *A. thaliana* and *S. parvula* exposed to different treatments for 24 hours. Cell wall yield was calculated as the percentage of plant biomass on a dry weight basis. In panels D-G, all values are mean ± SD (n=3~5). Asterisks represent significant differences (p<0.05) compared to control determined by Student’s t-test.

Next, we expanded our analyses of boron stress-responsive DEGs in functionally enriched clusters to all genes known to be involved in the biosynthesis of major cell wall components (Supplemental Data Set 1). We included both root and shoot tissues to view cell wall biosynthesis-related changes at the whole plant level. Figure 3B summarizes the cell wall biosynthesis pathways, including precursors, intermediates, and the building blocks of cell wall components, together with the genes involved in the process. Cellulose, hemicellulose, and pectin constitute about 90% of cell wall mass (Albersheim et al., 1996; Held et al., 2015). Cellulose is synthesized at the plasma membrane by cellulose synthase (CesA) complexes (Schneider et al., 2016; McFarlane et al., 2014), whereas pectins and hemicelluloses are assembled in the Golgi and then exported to the apoplast (Kousar et al., 2012). Hemicelluloses include xyloglucans, xylans, mannans, glucomannans, and mixed-linkage glucans (MLG) (Scheller and Ulvskov, 2010; Pauly et al., 2013), while the predominant pectins are homogalacturonan (HG), rhamnogalacturonan I (RG-I), and rhamnogalacturonan II (RG-II) (Atmodjo et al., 2013).

We found that a total of 14 out of 25 *galacturonosyltransferases* (*GAUTs*) and *GAUT-like* (*GATL*) genes, which were shown or suggested to be involved in pectin biosynthesis, were DEGs in *A. thaliana* shoots and roots, of which 13 were induced in shoots (Figure 3B, Supplemental Data Set 1). All the *glycosyltransferases* that have been proven or suggested to code for genes (4 *RGXTs* and 2 *SIAs*) involved in assembling the side chains of RG-II on a HG backbone showed a 4-fold or higher increase in *A. thaliana* shoots (Figure 3B, Supplemental Data Set 1). Those *glycosyltransferases* in the roots, however, did not show this induction (Figure 3B). Synergistically, several genes coding for the enzymes (UXSs and UXEs) producing UDP-arabinopyranose (UDP-Ara*p*) and UDP-arabinofuranose (UDP-Ara*f*), which are the donors used by glycosyltransferases to incorporate Ara*p* and Ara*f* into RG-I and RG-II (Bar-Peled and O’Neill, 2011), were also induced in *A. thaliana* shoots, while the majority of other NDP-sugar biosynthesis genes were repressed by excess boron (Figure 3B, left panels). Contrasting to the transcriptomic signal suggesting increased pectin components, the expression of all 10 *CesA* genes coding for cellulose synthases (Carroll and Specht, 2011; McFarlane et al., 2014) together with other genes mostly associated with hemicellulose biosynthesis remained unaffected (Figure 3B). The only experimentally verified molecular function of boron in plants is to cross-link RG-II-pectins in the cell wall (Kobayashi et al., 1996; Ishii et al., 1999; O’Neill, 2001; Funakawa and Miwa, 2015). Therefore, a net induction of genes associated with pectin biosynthesis in *A. thaliana* shoots suggested a path to potentially bind more boron and trap excess boron in shoot cell walls.

The cell wall also contains structural hydroxyproline-rich glycoproteins (HRGPs), notably the extensins (Cannon et al., 2008; Lamport et al., 2011). In *A. thaliana* shoots, excess boron significantly induced genes coding for multiple extensins and glycosyltransferases (GTs) involved in the extensin glycosylation (Velasquez et al., 2011) (Figure 3B and Supplemental Figure 4). It is notable that *A. thaliana* root and shoot expression profiles differed substantially. In roots, virtually all the transcripts potentially coding for extensins that significantly responded to excess boron were suppressed whereas in shoots, transcripts coding for extensins and associated GTs were all induced (Figure 3B). Such a differential regulation of the extensins suggests that the shoot cell wall may be stiffer in *A. thaliana* under excess boron than in the control (Supplemental Figure 4). Stiffening of cell walls has been reported in plants under other abiotic stresses (Tenhaken, 2015). In line with this finding, we also noted the co-repression of many of the genes encoding catalysts of cell wall loosening, including expansins and xyloglucan endotransglucosylase/hydrolases (XTHs) (Cosgrove, 2016) in both shoots and roots (Figure 3B, Supplemental Figure 4). The prominent changes in transcriptional responses related to cell wall biology observed in *A. thaliana* led us to hypothesize that cell walls serves as a sink to store excess boron under boron stress and the associated cell wall modifications were initiated as a transcriptional cascade of several processes including cell wall organization; synthesis and regulation of nucleotide sugar transporters that are linked to cell wall sugars; structural glycoproteins; and cell wall interacting kinases (Figure 3 A-B).

### Cell wall pectin precursor abundance significantly increased in response to excess boron in *A. thaliana* shoots

Based on the transcriptomic signature that suggested major cell wall modifications, especially related to pectins in *A. thaliana* shoots, we next assessed if such modifications coincided with changes in metabolic pools in boron-stressed shoots. We used GC-MS metabolic profiling to capture the primary metabolite pools from control and treated tissues of *A. thaliana* and *S. parvula*. We detected 39 and 70 annotated metabolites that were differentially accumulated under 5 mM and 10 mM boric acid treatments, respectively (Supplemental Data Set 3). By contrast, the relative abundances of any of these metabolites did not change significantly in *S. parvula* shoots exposed to excess boron (Supplemental Data Set 3).

We grouped the functionally annotated organic metabolites that significantly changed in their abundance in response to excess boron into five categories (Supplemental Data Set 3). At least one third of these metabolites were sugars or sugar derivatives, including sugar alcohols, while the remaining pool primarily consisted of amino acids and other amines, fatty acids, and other organic acids (Figure 3C). The organic acids included pyruvic acid, citric acid, and succinic acid. Their abundances are known to change in response to abiotic stresses, which is often associated with changes to overall energy balance during stress (Treves et al., 2020).

In line with our transcriptomic data, the relative abundance of many of the free monosaccharides that are components of cell wall polysaccharides, including arabinose, galactose, rhamnose, xylose, and mannose significantly increased upon excess boron in *A. thaliana* shoots (Figure 3D and Supplemental Figure 5). Remarkably, the abundance of none of these sugars significantly changed in boron-stressed *S. parvula* (Figure 3D). Arabinose, which is a component of RG-I and RG-II pectic polysaccharides (Bar-Peled and O’Neill, 2011), increased >2.5-fold, together with xylose, the precursor of arabinose (Atmodjo et al., 2013; Seifert, 2018) in response to excess boron in *A. thaliana* shoots (Figure 3D). Rhamnose, which is present in the side chains of RG-II and the backbone of RG-I (Bar-Peled and O’Neill, 2011), also showed a significant increase in response to 10 mM boric acid in *A. thaliana* shoots, together with galactose, another component of RG-I and RG-II. Similarly, the precursors of other cell wall polysaccharides, including mannose, fructose, and glucose increased in response to 10 mM boric acid (Figure 3D). Remarkably, none of these sugars changed significantly in *S. parvula* shoots during any of the boron treatments (Figure 3D). The overall changes in free monosaccharides during excess boron treatment of *A. thaliana* shoots support the view that pectic polysaccharides provide binding sites to trap excess boron in the cell walls.

We found that several sugar alcohols in *A. thaliana* shoots, including myo-inositol, cellobiotol, galactinol, erythritol, and glycerol, also increased in response to excess boron (Figure 3E). Myo-inositol is a precursor of pectin and hemicellulose (Kanter et al., 2005; Endres and Tenhaken, 2009), and its increase is consistent with our working model that cell wall pectins capture excess boron during boron stress. Cellobiotol and galactinol are also presumed to be involved in cell wall carbohydrate metabolism (Unda et al., 2017). Moreover, some of these sugar alcohols could bind excess boron in a manner similar to sorbitol and mannitol (Brown and Hu, 1998, 1996; Brown et al., 1999). Taken together, our metabolic profiling provide evidence that excess boron led to the increase in sugars and sugar alcohols, many of which are either directly or indirectly related to cell wall polysaccharides in *A. thaliana*.

### Boron accumulates in the cell walls of *A. thaliana* and *S. parvula*

We next determined if *A. thaliana* and *S. parvula* accumulated boron in their cell walls when grown in excess boron using ICP-MS. In *A. thaliana*, cell wall boron increased in roots and shoots of treated plants compared to the control group (Figure 3F). By contrast, we only detected an increase in boron in *S. parvula* root cell walls (Figure 3F). Cell wall yield remained constant under all tested conditions (Figure 3G), even 5 days after the treatments (Supplemental Figure 6). We performed a second series of experiments with four additional biological replicates using an extensive digestion procedure while minimizing possible contaminating boron sources during the experiment (see Methods) to validate boron sequestration in the cell wall during excess boron treatment. These results were consistent with the cell wall boron quantifications we initially performed (Pearson correlation coefficient *r* = 0.76, P = 0.015) (Supplemental Figure 7), and confirmed that boron accumulated in the cell walls of *A. thaliana* and *S. parvula*.

We next determined if other elements present in plant tissues accumulated in the cell walls as a result of excess boron treatments. None of the 20 elements analyzed differed significantly from control plants in cell walls from *S. parvula* roots and shoots and *A. thaliana* shoots (Supplemental Figure 8). By contrast, almost half of the elements decreased in abundance in root cell walls of excess boron-treated *A. thaliana* (Supplemental Figure 8). This is likely related to the substantial root growth inhibition observed specifically for excess boron-treated *A. thaliana*.

Our results, when taken together, suggest that cell walls do capture excess boron. Additionally, *A. thaliana* shoot cell walls have a higher capacity to retain boron than their root counterparts (Figure 3F). Since cell wall yield did not change in response to excess boron, the observed changes are likely due to alterations in the internal structures of the cell walls to enable compartmentalization of excess boron. We also observed that the increase in the boron content in *A. thaliana* shoot and root cell walls (>2 fold) was smaller than the increase in boron content in the entire tissue (compare Figures 1H and 3F). This was most notable in *A. thaliana* roots where whole-tissue boron content increased up to 27 fold. A less pronounced, but similar trend was observed for *S. parvula* roots (Figure 3F). Therefore, cell walls may only provide a partial sink for excess boron, and cellular processes involved in cell wall modifications may be limited in the amounts of boron that they can sequester.

### Altered RNA metabolism in response to excess boron led to an increased mean expression of the entire transcriptome in *A. thaliana* roots and shoots

We next searched for cellular processes that may serve as additional substrates for excess boron or molecular targets that may cause cellular toxicity if bound by excess boron in the cytoplasm. We expected that the genes induced by excess boron may shed light on such processes. Our analysis identified 19 functional clusters comprised of 2,112 of the 2,622 boron stress-induced genes in *A. thaliana* roots, of which the top ten largest clusters represented 99% (2,094) of the total number of genes from all clusters (Supplemental Data Set 2). The most notable cellular process among the induced genes in roots was RNA metabolism described by the largest functional cluster (C1 in Figure 4A). Further, clusters C5, included transcription and translation regulation, and C9, represented by ribosome organization, are also associated with RNA metabolism (Figure 4A, red boxes). This indicates that in *A. thaliana* roots, RNA metabolism-related processes were substantially affected by boron toxicity.

**Figure 4.**
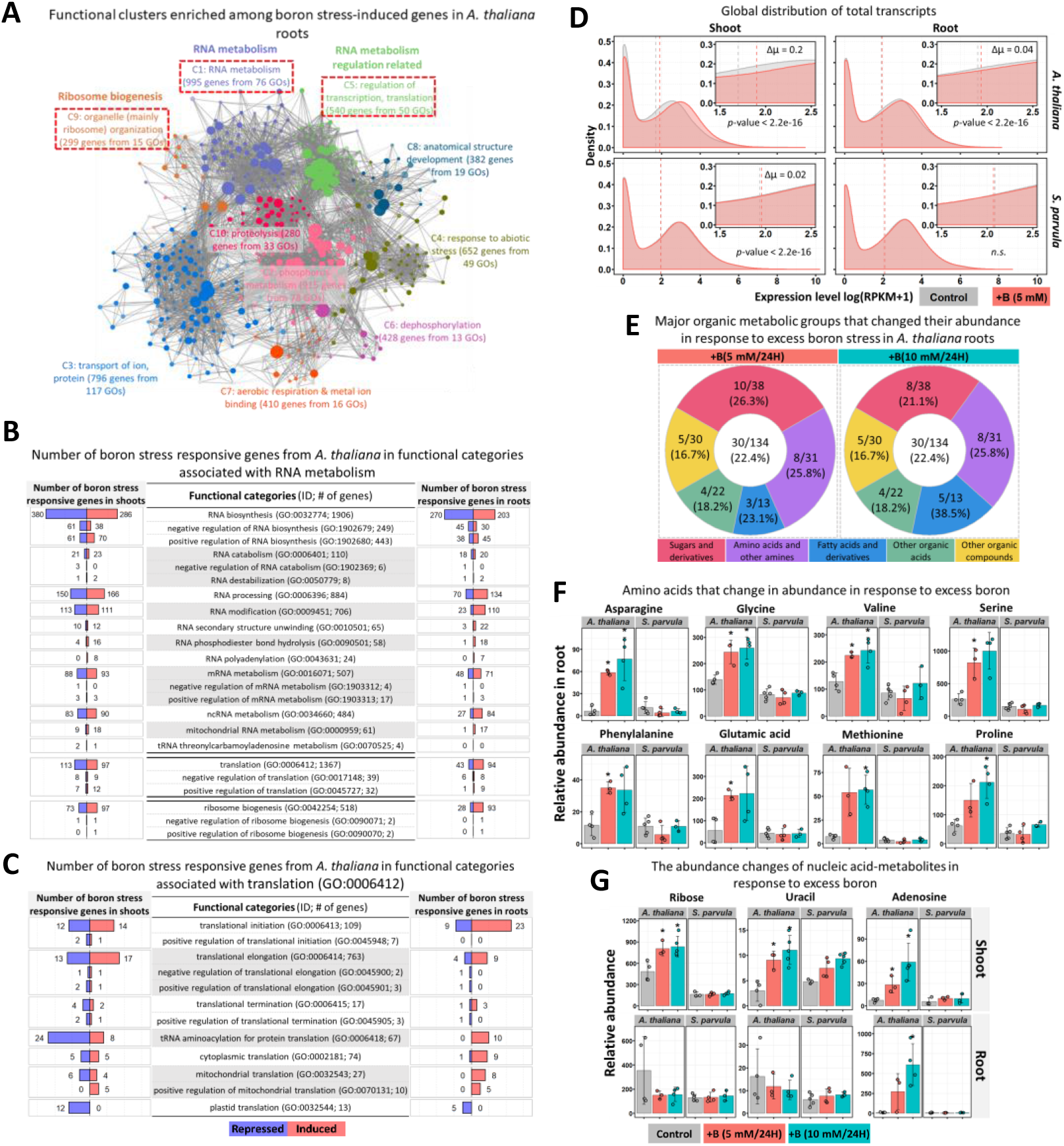
RNA metabolism related processes are affected in response to excess boron in *A. thaliana*. (A) Enriched functional clusters among boron stress-induced genes in *A. thaliana* roots with notable associations for RNA metabolism marked by red-dashed boxes. The network visualization is similar to Figure 3A. (B-C) Number of boron stress responsive genes from *A. thaliana* in functional categories associated with RNA metabolism (B) and translation (GO:0006412). (C). Red and blue indicate up- and down-regulation, respectively. (D) Global transcriptome expression distributions in *A. thaliana* and *S. parvula*. Density distributions of expression in log(RPKM+1) for entire transcriptomes were compared to identify transcriptome-wide global changes. *p*-values were estimated using Wilcoxon signed-rank test. (E) Major organic metabolic groups that changed their abundance in response to excess boron stress in *A. thaliana* roots. For each category, the number and proportion of metabolites that changed in abundance compared to the total identified are shown. (F) Amino acid and (G) nucleic acid-metabolite abundance changes in response to excess boron stress in shoot and root. The relative abundance is given compared to the internal standard, ribitol. Values shown are mean ± SD (n = 3, 4 or 5). Asterisks represent significant differences of each treatment compared to control according to Student’s t test (p < 0.05).

To further investigate how RNA metabolism could be altered by excess boron, we examined all genes represented by RNA metabolism (GO:0016070), together with their regulators annotated under specific child GO terms in the *A. thaliana* genome. Interestingly, all RNA metabolism processes, as well as translation and ribosome biogenesis, were enriched in genes differentially responsive to excess boron in our study (Figure 4B). This further confirmed our earlier observation of RNA metabolism being a major target of boron stress, especially in *A. thaliana* roots (Figure 4A). There were many more boron stress-induced genes than repressed genes in most of the GO categories especially in *A. thaliana* roots (Figure 4B). For example, RNA processing, RNA modification, ncRNA metabolism, RNA metabolism, RNA secondary structure unwinding, RNA polyadenylation, translation, and ribosome biogenesis all had more genes induced than repressed in each category in roots (Figure 4B, 4C).

If RNA metabolism was the most dominant process among the boron stress-induced genes in *A. thaliana*, we hypothesized that the stress effect should be discernible at the entire transcriptome level. Therefore, we tested if the mean expression level per transcript for the entire transcriptome was significantly shifted in the excess boron-treated samples compared to the control, as previously described by Muyle and Gaut (2018). Excess boron stress did lead to an increased mean expression in *A. thaliana* roots and shoots and also in *S. parvula* shoots (Figure 4D). The boron stress-adapted *S. parvula*, however, did not show a mean expression change in roots implying its greater capacity to cope with excess boron without a massive change to its entire transcriptome. This may also suggest that the Arabidopsis global transcriptomic response has a significant energetic cost, which could also contribute to the delayed growth during excess boric acid treatments observed (Figure 1).

### *A. thaliana* roots respond to excess boron by increasing the abundance of multiple amino acids, sugars, and nucleic acid-metabolites

We observed induction of genes especially associated with translation (Figure 4C) and ribosome biogenesis (Figure 4B) in *A. thaliana* roots, which was further supported by the increased average expression level observed for the entire *A. thaliana* root transcriptomes (Figure 4D). This led us to test whether changes in RNA metabolism and associated processes observed in the root transcriptomes of *A. thaliana* during excess boron stress affected amino acid usage. Additionally, we suspected that an increased level of translation may also lead to altered metabolic pools of sugars involved in primary energy metabolism especially in *A. thaliana* roots in response to excess boron.

The abundance of 32 functionally annotated metabolites changed significantly upon 5 or 10 mM boron treatments in *A. thaliana* roots (Supplemental Data Set 3). In contrast, none of these metabolites were affected by either boric acid treatment in *S. parvula* roots (Supplemental Data Set 3). Sugars and amino acids, and their derivatives, constituted the two largest groups of boron stress-responsive metabolites in *A. thaliana* roots (Figure 4E). The relative abundance of 8 of the 13 amino acids detected significantly increased in the root tissues in response to the treatments (Figure 4F and Supplemental Data Set 3). This response was more prominent in roots than shoots, where only two out of the 10 amino acids detected changed significantly (Supplemental Data Set 3). The majority of sugars and their derivatives that responded to the treatments in the roots were primarily involved in glycolysis or sugar transport. For example, these included glucopyranose, fructofuranose, glucose 1-phosphate, glucose 6-phosphate, fructose 6-phosphate, sucrose, and raffinose (Supplemental Data Set 3). This may be indicative of the generally higher demand for cellular energy consumption during induced transcription levels especially in *A. thaliana* roots under excess boron stress. Notably, the sugar-metabolite profile in the roots (Supplemental Data Set 3) was quite distinct from the increased abundance of sugars in the shoots that were enriched primarily in cell wall precursors as described earlier (Figure 3D). Ribose, uracil, and adenosine that are related to nucleic acid metabolism also increased in abundance in shoots in response to excess boron, whereas only adenosine from that group increased in the roots (Figure 4G). Adenosine was the metabolite with the highest fold change in shoots and roots among all metabolites detected in our study (Figure 4G).

### Constitutive expression of *S. parvula* orthologs match the post-stress expression of *A. thaliana* boron stress-responsive orthologs

To complement our studies focused on boron stress-sensitive *A. thaliana*, we next sought evidence for the types of biological processes that allow *S. parvula* to tolerate toxic amounts of boron. The overall transcriptomic, ionomic, and metabolomic responses elicited in *S. parvula* in response to excess boron were much less pronounced than those for *A. thaliana*. Nevertheless, we were able to observe enriched functions among the differentially expressed genes (Figure 5). Cell wall-modifying enzymes were the only enriched function observed for *S. parvula* roots (Figure 5, Supplemental Data Set 2). Genes encoding protein modifying and mitochondria-localized proteins were also induced in response to the boric acid treatments in *S. parvula* shoots, while genes involved in biotic stress and defense responses and boron uptake were repressed in both roots and shoots (Figure 5, Supplemental Data Set 2).

**Figure 5.**
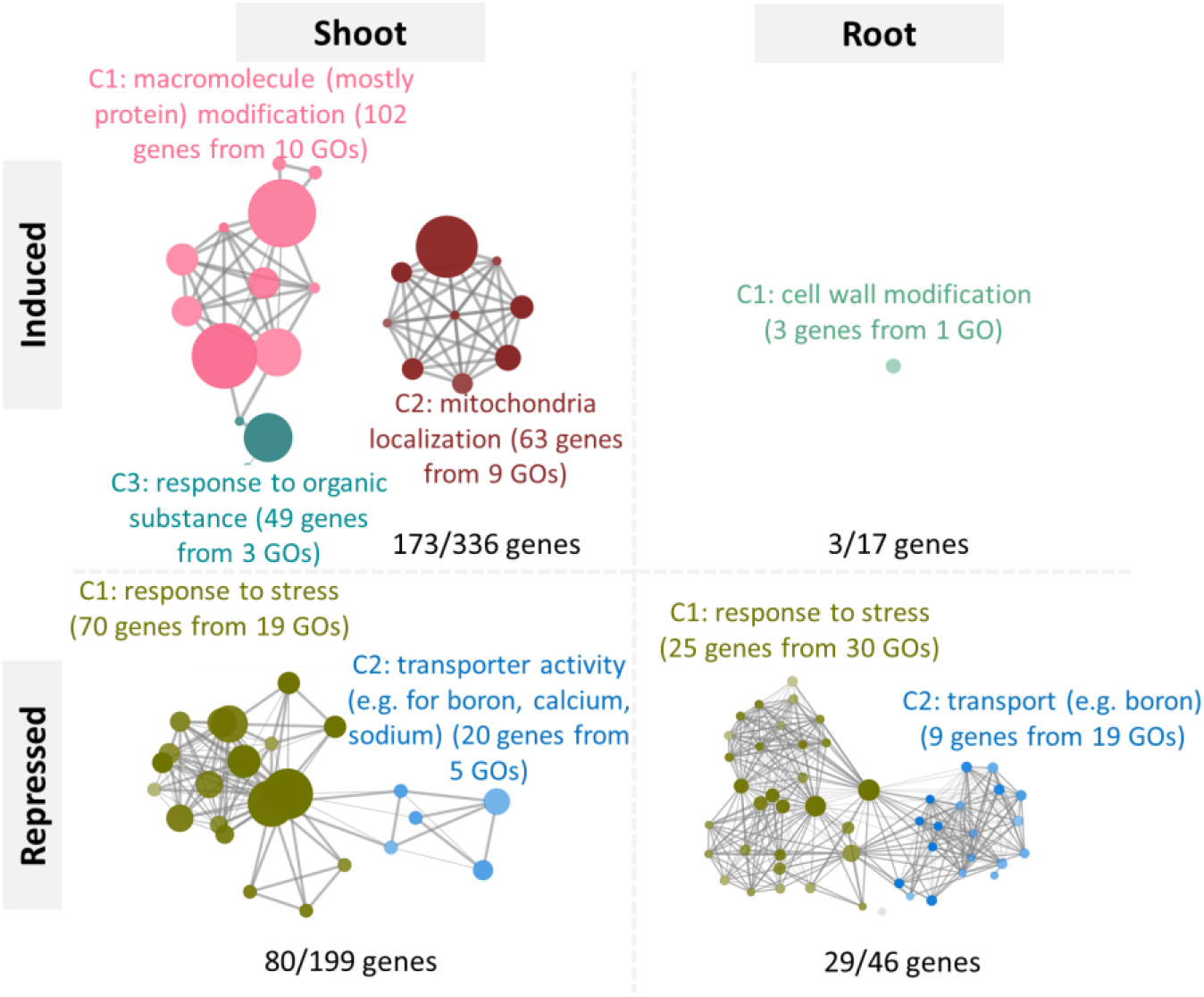
Functional clusters enriched among differentially expressed genes in *S. parvula* in response to excess boron. The network visualization was constructed the same way described for Figure 3A. Numbers assigned for each cluster represent total number of genes from all subclusters/total number of boron stress-responsive genes in the category.

Our comparative -omics framework allows us to gain insight into the *S. parvula* genes and processes that remained unchanged when their orthologs were differentially regulated in *A. thaliana* in response to excess boron. To this end, we compared the expression levels of orthologs from the two species under control and treated conditions. We identified ortholog expression for 19,263 pairs in shoots and 19,784 pairs in roots that were used to determine the co-expressed ortholog clusters. This led to 22 shoot and 20 root clusters (Supplemental Data Set 4). We further categorized those clusters into four overall expression trends that we have termed as, (a) stress-ready clusters; (b) unique-response clusters; (c) shared-response clusters; and (d) no-response clusters (Figure 6A). In the stress-ready cluster (a), an ortholog from one species responded to the stress to reach a level of expression equivalent to the basal level of the ortholog in the other plants, which itself remain unchanged under the stress. The unique-response clusters (b) represented ortholog pairs where one species showed a response that was unmatched in the other species either at the control or treated levels. Orthologs pairs with a similar response in both species to excess boron stress were categorized into the shared-response cluster (c). Finally, ortholog pairs that did not change their expression to excess boron stress were grouped as “no response” (d).

**Figure 6.**
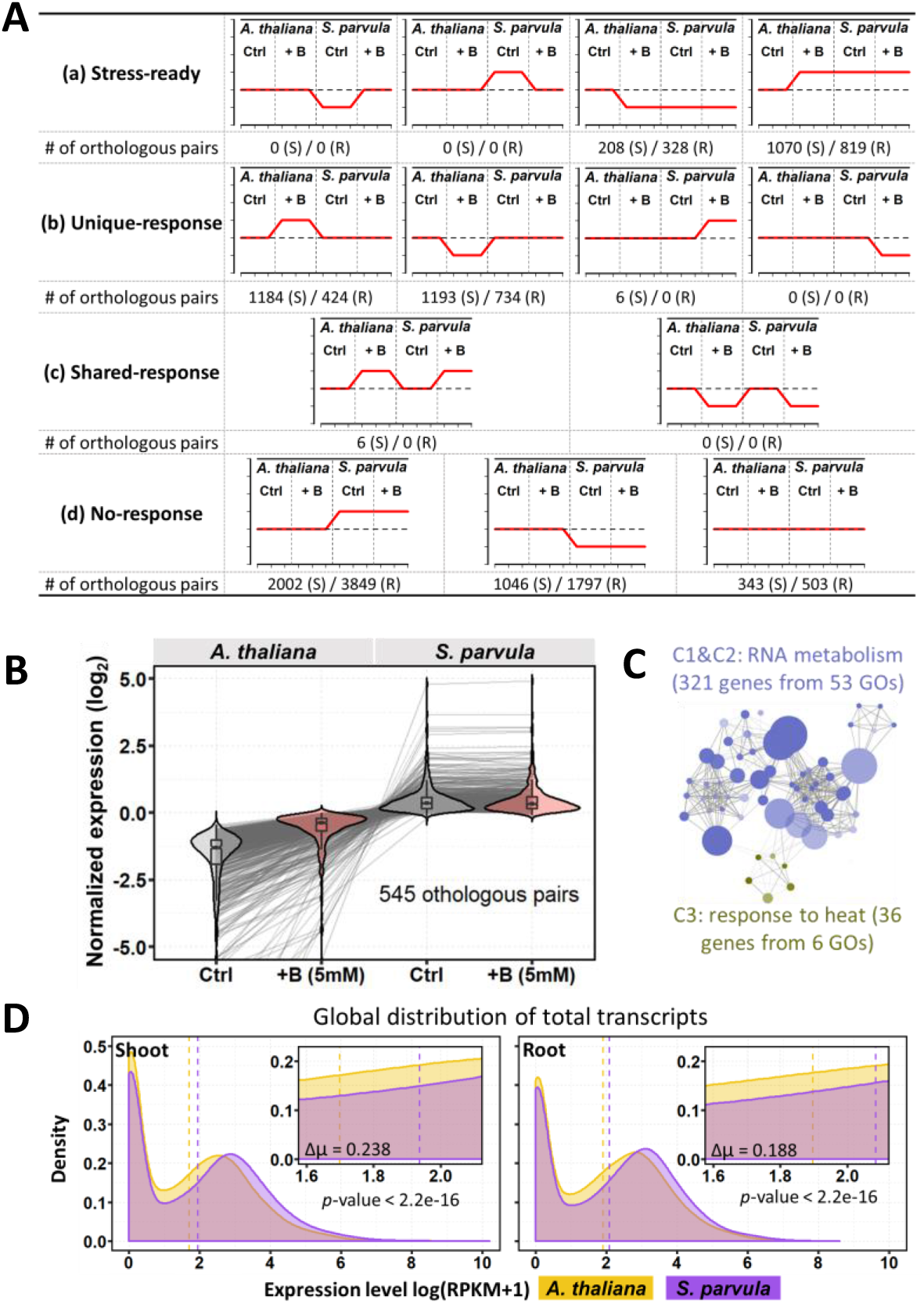
Genes associated with excess boron responses are constitutively expressed in *S. parvula*. (A) Summary of co-expression trends in ortholog pairs between *A. thaliana* and *S. parvula* in response to excess boron. The red lines indicate the trend in expression levels of the ortholog pairs in each species under control and treatment conditions, compared to *A. thaliana* control (dashed line). Ctrl, Control; + B, boric acid treatment; S, Shoot; R, Root. (B) A major co-expression cluster in the “stress-ready” category in (A) that illustrates stress preparedness of the *S. parvula* orthologs in roots (center line, median; box, interquartile range (IQR); notch, 1.58 × IQR / sqrt(n); whiskers, 1.5 × IQR). (C) Functional clusters enriched among the ortholog pairs from the example cluster given in (B). The clusters are differently colored and labelled with the representative functions. Node size represents genes in the test set which are annotated to that functional term; edges represent the number of shared genes between functional terms; each cluster is coded with a different color; and shade of each node represents p-value assigned by the enrichment test. Lighter to darker shades indicate larger to smaller *p*-values, respectively. (D) *S. parvula* shows a higher basal level expression than *A. thaliana*. *p*-values were estimated using Wilcoxon signed-rank test.

The majority of the ortholog pairs in the two species remained unchanged in response to excess boron (7,797 ortholog pairs from roots and shoots in the no-response group) (Figure 6A). When one ortholog in a pair did respond, the majority (~62%) of those showed only the expression change in the *A. thaliana* ortholog. The ortholog distribution in these categories further highlighted the more restrained transcriptomic responses of *S. parvula* and revealed an interesting but hidden feature of the *S. parvula* genome that we may not have identified without *A. thaliana* as a comparator. We did not identify a single ortholog pair where the *S. parvula* ortholog responded to the stress to reach the basal level of its *A. thaliana* ortholog (i.e. zero representation in the stress-ready group for *A. thaliana*). By contrast, we identified 2,160 *A. thaliana* orthologs whose expression changed to match the basal expression observed for the *S. parvula* orthologs. Additionally, we only identified 6 *S. parvula* orthologs that could be classified in the unique-response group (Figure 6A). This led us to propose that stress-adapted *S. parvula* had a pre-adapted transcriptome with over a thousand orthologs whose basal expression levels (pre-stress expression) match the expression levels achieved in response to the stress (post-stress expression) in stress-sensitive *A. thaliana*. Any differential expression shown by the orthologs in the stress-adapted species prompted by the stress was always echoed by the stress-sensitive species. Thus, these cellular responses may be common among plants responding to excess boron and not restricted by species boundaries.

We also identified at least one thousand orthologs in *A. thaliana* roots and shoots that uniquely responded to excess boron. The expression of their *S. parvula* counterparts did not change significantly. We suspect that the majority of the expression changes in *A. thaliana* represent non-specific symptoms caused by interruption to cellular processes in a plant unable to sustain a cellular environment conducive for growth and development rather than a specific response to excess boron.

We also searched for enriched functions associated with the stress-ready clusters in *S. parvula* to determine what cellular or metabolic processes were enriched at stress-anticipatory levels in the basal transcriptomes. We first looked into the orthologs expressed in the *S. parvula* stress-ready category where *A. thaliana* orthologs were induced in response to excess boron (Figure 6B). These *S. parvula* orthologs were predominantly enriched for RNA metabolic processes (Figure 6C). It should be noted that the same enriched function was also the predominant function among all induced genes in *A. thaliana* roots regardless of their orthologous relationship with *S. parvula* (Figure 4A). We then compared the basal expression levels of all orthologs between *A. thaliana* and *S. parvula* to assess if the basal expression was significantly different between the two species. In both shoots and roots, the *S. parvula* transcriptome showed significant shifts towards overall higher gene expression levels compared to *A. thaliana* (Figure 6D). Taken together, these results suggested that *S. parvula* transcriptomes were pre-adapted for boron stress most notably in the metabolic functions associated with RNA metabolism that was among the most altered processes in the stress-sensitive *A. thaliana* during the excess boron treatments. However, in many of these stress-ready clusters, enriched functions only described a subset of the orthologs, while a significant proportion of orthologs remained functionally uncharacterized (Supplemental Figure 9).

## Discussion

Combining our results and previous studies, we propose a model for how excess boron triggers transcriptomic responses that cascade into major cellular and growth responses (Figure 7). The stress-sensitive species, *A. thaliana*, in response to boron toxicity: 1) halts active boron uptake; 2) deposits a proportion of excess boron into cell walls; 3) adjusts the expression of genes involved in RNA metabolism; and 4) forms complexes with free boric acid, especially in roots. We demonstrated that boron toxicity induced minimal changes to gene expression, elemental and metabolite profiles, and growth in stress-adapted *S. parvula* when compared to *A. thaliana*. Different excess boron tolerance mechanisms are likely present in *S. parvula.* These include, 1) an efficient boron efflux system that minimizes excess boron accumulation in the plant; 2) cell wall absorption of a proportion of excess boron; 3) formation of B-complexes to reduce free boric acid accumulated in the cytoplasm before boron could bind to essential metabolites; and 4) genes associated with cellular processes affected by excess boron in *A. thaliana* are constitutively expressed at stress pre-adapted levels.

**Figure 7.**
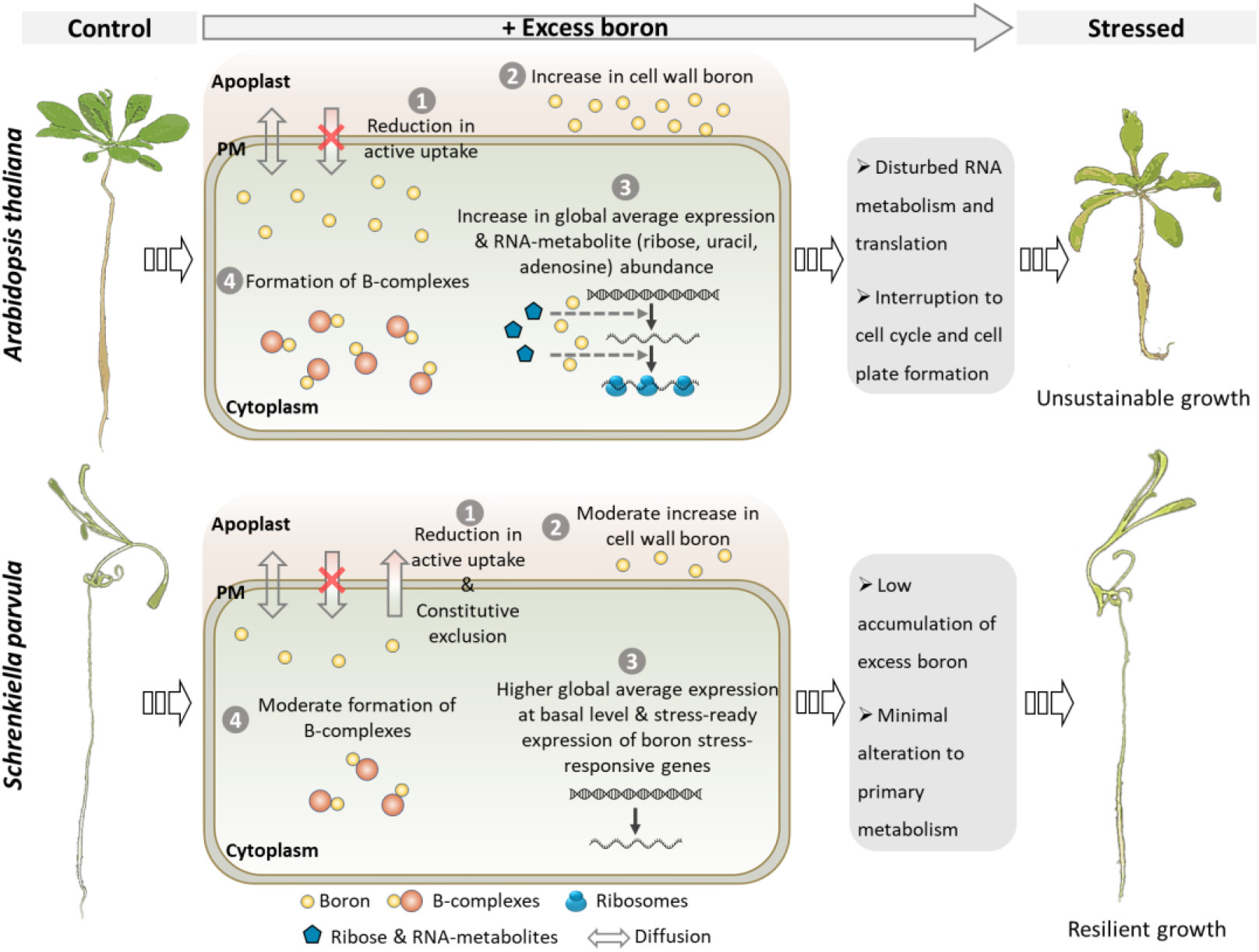
A proposed model for boron toxicity in plants.

### *S. parvula* is equipped with an efficient boron efflux system

*S. parvula* is an extremophyte that has evolved to grow on boron-rich soils (Nilhan et al., 2008; Oh et al., 2014). As expected, it was less affected by excess boron than the boron stress-sensitive model, *A. thaliana* (Figure 1). This is due in part to the ability of *S. parvula* to maintain relatively low boron levels in its tissues (Figure 1H). This is likely a feature of boron toxicity tolerance since other boron stress-tolerant plants, including *Eutrema salsugineum* (Lamdan et al., 2012) and *Puccinellia distans* (Stiles et al., 2010), also maintain a relatively low level of endogenous boron even when grown under excess boron conditions.

At physiological pH, boron primarily exists as uncharged boric acid, which is highly membrane permeable (Reid, 2014). Boric acid readily diffuses into the root cells under adequate or excess boron conditions (Yoshinari and Takano, 2017; Landi et al., 2019; Princi et al., 2016). Several mechanisms have evolved in plants to control boron influx and efflux. For example, *A. thaliana BOR4* encodes the only boron exporter experimentally shown to function under boron toxicity (Miwa et al., 2014, 2007). Surprisingly, we saw no significant change of expression of this gene in either species in response to excess boron. However, *BOR5*, the closest homolog of *BOR4*, was induced by excess boron in *A. thaliana* roots, and was highly expressed especially in the roots of *S. parvula* control plants (Figure 2C, D). This may be a result of a 15 kb transposition insertion in the upstream region adjacent to the *SpBOR5* transcription start site (Oh et al., 2014). *SpBOR5* and *AtBOR5* exist as single copy genes and are co-linear except for the genomic insertion in *S. parvula* (Oh et al., 2014; Oh and Dassanayake, 2019). We demonstrated that *SpBOR5* is an effective boron exporter (Figure 2D) and propose that it is likely a key contributor to the underlying tolerance of *S. parvula* to excess boron (Figure 7).

### Excess boron taken into plants is differently compartmentalized in *A. thaliana* and *S. parvula*

The absorbed excess boron may exist in free or bound forms in plants. We observed that free boric acid levels increased in *A. thaliana* shoots and roots, as well as in *S. parvula* shoots as the external boric acid concentration increased (Figure 1H). Plants may attempt to minimize the deleterious effects of excess boric acid by exporting it to vacuoles. However, we saw no change in the expression of *TIP5;1*, which encodes the only known aquaporin that facilitates boron transport into vacuoles (Pang et al., 2010), in either *A. thaliana* or *S. parvula* (Supplemental Figure 10A). Other boron stress-responsive TIP genes all showed repression instead of induction in treated *A. thaliana* (Supplemental Data Set 1).

In a previous study of two barley cultivars that differed in their boron tolerance, the boron stress-tolerant cultivar was reported to have a higher apoplastic boron content than in the sensitive cultivar (Reid and Fitzpatrick, 2009a). We found that the expression levels of *AtBOR5* and *AtBOR7* increased in *A. thaliana* roots in response to excess boric acid (Figure 2C and Supplemental Data Set 1). Therefore, it is possible that a proportion of excess free boric acid is exported into the apoplast, especially in *A. thaliana* roots. Consistent with this notion, previous studies have suggested that apoplastic boric acid constitutes the majority of soluble boron in plants under normal conditions, and even in some species after exposure to excess boron (Matoh, 1997). The increase in free boric acid in *A. thaliana* and *S. parvula* is unlikely to be the sole cause of the increased amounts of total boron detected (Figure 1H and I). Rather, some absorbed boron must exist in a bound form especially in *A. thaliana* roots and *S. parvula* shoots. The formation of B-complexes may have contributed to the detoxification of excess boron. Alternatively, such complexes may also accumulate in the cytoplasm as undesirable metabolic end products.

Our metabolomic profiles indicated that ribose increased in *A. thaliana* shoots under excess boron (Figure 4G). This monosaccharide together with ribose-containing compounds, including nucleotides, NADH, NAD^+^, and S-adenosylmethionine have the ability to form borate esters in the cytoplasm (Ricardo, 2004; Ralston and Hunt, 2001; Kim et al., 2003, 2004). It is notable that adenosine is among the largest metabolite changes (~65 fold increase) in treated *A. thaliana* (Figure 4G). Boron could also form borate esters with sugar alcohols and organic acids containing *cis*-diols (Bolanos et al., 2004). Several sugar alcohols, including galactinol, erythritol, and cellobiotol increased substantially in treated *A. thaliana* shoots (Figure 3E). We also observed that many unidentified compounds changed in *A. thaliana* during boron treatments. Remarkably, none of the identified metabolites changed significantly in *S. parvula* in response to excess boron treatments. This is consistent with our hypothesis of a transcriptome pre-adapted to boron stress in the tolerant *S. parvula* (Supplemental Data Set 3).

The lack of substantial changes in the metabolite profiles of *S. parvula* led us to hypothesize two possibilities for how it may minimize the cellular toxicity of excess boron in the cytoplasm. First, generation of borate-containing metabolites may ameliorate toxicity but comes with a high energy cost that would direct *S. parvula* to use more energy efficient alternative paths to store excess boron. Second, if the generation of such B-complexes was harmful but unavoidable when excess boron accumulated in the cytoplasm, *S. parvula* may prevent their accumulation by limiting the amounts of boron in the cytoplasm more efficiently than *A. thaliana*. When bound to boron, metabolites in the cytoplasm will be unavailable to critical primary metabolic processes. Thus, cells may attempt to increase the production of these metabolites at a rate that cannot be sustained in boron stress-sensitive species. The response of *A. thaliana* to increase many of these metabolites on excess boron are consistent with this view. Alternatively, mechanisms may have developed in *S. parvula* to process excess cytoplasmic boron in a manner that does not preclude ribose or other metabolite pools from functioning in their respective essential roles (Figure 7).

### Cell wall contributes to partially compartmentalize excess boron

Several independent studies have provided compelling evidence for the existence of boron-rhamnogalacturonan-II (B-RG-II) complexes in plant cell walls (Kobayashi et al., 1996; Ishii and Matsunaga, 1996; O’Neill et al., 1996). There is also evidence that this complex is required for normal plant growth and development (Fleischer et al., 1999; Ishii et al., 2001; O’Neill, 2001). The carbohydrate–rich plant cell wall is ideally suited to bind boron (Matoh, 1997), but whether cell walls can store excess boron when plants encounter boron toxicity has not been demonstrated. Herein, we provide compelling evidence for this phenomenon. First, we found that while cell wall yield was unaffected, there was an increase in cell wall boron in *A. thaliana* shoots and roots, as well as in *S. parvula* roots, when plants were grown on excess boron (Figure 3F, G). Second, we have demonstrated that boron toxicity altered the expression of many genes involved in cell wall biogenesis or organization as well as pectin biosynthesis (Figure 3A, B). Third, our metabolomic profiling supported the transcriptomic signals related to the changes in the content of cell wall polysaccharide precursors, notably the monosaccharides used to synthesize pectin (Figure 3D). Together these observations strongly support the idea that cell walls contribute, at least partially, to the compartmentation of excess boron in plants (Figure 7).

In line with our results, previous studies on *A. thaliana* and boron stress sensitive citrus cultivars showed boron accumulation in the cell sap-free tissue fraction when treated with excess boron (Lamdan et al., 2012)(Martínez-Cuenca et al., 2015). A recent study of the trifoliate orange (*Poncirus trifoliata*) reported alterations in cell wall structure when plants were treated with excess boron (Riaz et al., 2019; Wu et al., 2019). In contrast to these findings, Dannel et al. (1998) suggested that cell walls did not absorb excess boron during boron toxicity based on studies of boron stress-resistant sunflowers. However, they did not quantify boron accumulation in tissues, and assumed that internal boron levels changed proportionally to the external boron supply; thereby ignoring the possible contribution of active extrusion of excess boron in plants. A subsequent study reexamined boron tolerance in sunflower and concluded that sunflower did exclude excess boron when compared to a sensitive species (Keleş et al., 2011). Several other studies, for example, have noted that barley roots (Hayes and Reid, 2004) and *Eutrema salsugineum* shoots (Lamdan et al., 2012) did not store excess boron in the corresponding cell walls. However, these studies did not include both roots and shoots when assessing how excess boron could be partly stored in certain tissues while some of it could be extruded back to the soil.

In cell walls, boron can complex with apiose present in RG-II as well as with other sugars containing *cis*-diols (Matoh, 1997). Boron cross-linking of two RG-II molecules occurs rapidly during RG-II synthesis and secretion. Previous studies suggest that the crosslink is formed in the cytoplasm prior to RG-II deposition in cell wall rather than in the cell wall itself (Chormova et al., 2014; Chormova and Fry, 2016). *In vitro* assays have demonstrated that excess boron can reduce the rate of RG-II dimerization (Chormova et al., 2014). Therefore, future studies testing the compositional changes of RG-II and other cell wall sugars during excess boron stress in plants could further identify how plant cell walls may be restructured to allow storage of excess boron.

### Boron toxicity disturbs RNA metabolism and related processes

Excess boron resulted in substantial changes in the expression of genes involved in RNA metabolism and related processes, including translation and ribosome biogenesis (Figure 4A). Boron is known to form complexes with ribose (Ricardo, 2004) and ribose-containing compounds *in vitro* (Ralston and Hunt, 2001; Kim et al., 2003, 2004). Thus, one explanation for the extensive changes in RNA metabolism-related processes could be that excess boron affects the availability of ribose and ribose-containing compounds needed for RNA metabolism, and that creates a prominent transcriptional footprint.

Uluisik *et al*., (2011) previously demonstrated that excess boron suppresses protein synthesis and interrupts translation initiation by reducing the proportion of functionally available polysomes in yeast. The authors further showed that excess boron also inhibits aminoacylation of tRNAs *in vitro*. Considering our transcriptomic and metabolomic results, together with the previous publications, it is reasonable to suspect that similar to yeast, excess boron in plants may impact protein synthesis by impairing polysome function. In addition, excess boron may also bind to the ribose moiety at the amino acid attachment site in tRNAs, which could block access to amino acids, thus inhibiting tRNA aminoacylation. In support of this view, our transcriptomic data shows that ribosome biogenesis was enhanced in *A. thaliana* roots and shoots after excess boron treatments (Figure 4B and 7).

### *S. parvula* transcriptome is pre-adapted to boron toxicity

Compared to *A. thaliana*, *S. parvula* is more tolerant to boron toxicity (Figure 1). Our transcriptomic analyses suggest that *S. parvula* is pre-adapted for this stress (Figure 6A). While some of the *S. parvula* orthologs in the “stress-ready” cluster could be readily associated with enriched GO functions (Figure 5), not all orthologs could be represented by GO annotations inferred using experimentally established functions (Supplemental Figure 9). The proteins encoded by many of these genes (>50% in stress-ready clusters) have no known functions described for their *A. thaliana* orthologs. This indicates a severe gap in the functional associations recognized between gene functions relevant to excess boron stress. Our comparative transcriptome analyses indicate that these genes of unknown functions in *A. thaliana* not only respond significantly to excess boron, but also their orthologs in *S. parvula* are expressed at levels comparable to the induced or repressed level in *A. thaliana* even in the absence of boron stress. Such stress-preparedness at the transcriptome level is likely a key contributor to the stress response in boron stress-tolerant plants. Indeed, similar transcriptome-level preadaptation to other abiotic stresses have been documented for plants that have evolved in environments where abiotic stresses are a constant feature (Taji et al., 2004; Gong et al., 2005; Becher et al., 2004; Hassan et al., 2016).

### Why is excess boron toxic to plants?

Our results demonstrated that when plants are grown in the presence of excess boron, some of this boron accumulates in cell walls. However, incorporating boron beyond an undefined threshold may trigger cell wall integrity signaling. We found >55% of genes (at least 300 in shoots and 150 in roots out of 628) coded for receptor-like kinases (RLKs) that responded to excess boron in *A. thaliana* (Supplemental Data Set 1). Many of these genes including *wall-associated kinases (WAKs), Catharanthus roseus RLK1 (CrRLK1)-like (CrRLK1L) kinases,* and *leucine-rich repeat (LRR) RLKs* have been suggested to participate in cell wall integrity sensing (Steinwand and Kieber, 2010; Rui and Dinneny, 2019; Vaahtera et al., 2019).

We observed that excess boron in *A. thaliana* shoots led to the repression of several *cellulose synthases*, including *CesA2* and *CesA3*, and *CesA* like family members (*CSLD5*) (Supplemental Figure 10B). *CSLD5* is most highly expressed in the shoot meristem of *A. thaliana* and is required for initializing cell plate formation (Gu et al., 2016). Boron-dependent repression of *CSLD5* may result in arresting cells in their G2/M transition phase, leading to cell division failures and growth defects. Further, excess boron is reported to decrease the number of mitotic cells and increase the fraction of 4C cells in *A. thaliana* root tips (Sakamoto et al., 2011). Additional studies have reported that inhibition of cellulose biosynthesis leads to the repression of cell cycle genes (Gigli-Bisceglia et al., 2018) and that key core cell cycle regulators are modulated by excess boron (Aquea *et al.* 2012). Our data are consistent with these publications, as we identified cell cycle processes together with exocytosis, which is related to cell-plate formation, as major functional groups among boron stress-repressed genes in *A. thaliana* shoots (Supplemental Figure 11). Therefore, excess boron accumulation in cell walls may not only affect cell wall integrity, but also cell plate construction, which in turn may interrupt cell division. This may explain why the effects of excess boron become apparent in fast-dividing meristems before mature tissue (Choi et al., 2007; Reid et al., 2004; Aquea et al., 2012).

Excess boron is not only toxic to plants, but also to yeast and animals (Bakar Salleh et al., 2010; Bakirdere et al., 2014). Therefore, cell wall-mediated boron toxicity alone may not explain the toxic effects of excess boron on these systems, especially animal cells. Excess boron-associated DNA damage has been reported as a consequence of boron toxicity among eukaryotes (Sakamoto et al., 2018). In addition, we showed that transcriptional signals related to RNA metabolism were substantially affected in *A. thaliana*, while *S. parvula* orthologs showed a stress-prepared expression level prior to the stress (Figure 4A, 6A, and 6B). We also observed transcriptome responses pointing to translation as a major target of boron toxicity. Similar results have been reported for yeast (Uluisik et al., 2011). Further, in human cells, excess boron increased the phosphorylation of eIF2α, which was inferred to lead to reduced protein synthesis (Yamada and Eckhert, 2018; Henderson et al., 2015).

In conclusion, we have shown that boron toxicity induces significant physiological and molecular changes in boron stress-sensitive *A. thaliana* compared to stress-adapted *S. parvula*. Excess boron accumulates in the cell walls of both shoots and roots, which may alter the structure and properties of the cell wall and its components. Such changes in the cell wall may affect cell plate formation, which in turn may lead to interruptions in cell division. Our data also suggest that boron toxicity interferes with RNA metabolism-related processes, especially translation, and other metabolic processes that involve ribose-containing metabolites. A model for how excess boron may trigger transcriptomic responses that cascade into major cellular and growth responses is presented in Figure 7. Further studies into cell wall dynamics during excess boron treatments in *A. thaliana*, as well as targeted functional analyses of *A. thaliana* stress-responsive genes that also show “stress-adapted” transcription in *S. parvula* to determine their currently unexplored functions would lead to an extended overview of how plants can survive excess boron stress.

## Materials and Methods

### Plant material and growth conditions

*Schrenkiella parvula* (ecotype Lake Tuz) and *Arabidopsis thaliana* (ecotype Col-0) seeds were surface sterilized with Clorox diluted 1:1 containing 0.05% Tween-20 and 70% ethanol, followed by 4-5 washes with sterile dH_2_O. Sterilized seeds were stratified for 4 days at 4 °C in the dark.

Plants for RNAseq, metabolomics, and ionomics experiments, were grown hydroponically in 1/5-strength Hoagland’s solution (Liu et al., 2010; Wang et al., 2018) at 22°C to 24°C in a growth chamber with a 14-h-light/10-h-dark cycle; 100-150 μmol m^−2^ s^−1^ light intensity. 4-week-old plants were transferred to growth media containing fresh 1/5-strength Hoagland’s solution or Hoagland’s solutions containing 5, 10, or 15 mM boric acid. The pH of the media was measured and adjusted to match control solutions in growth media allocated for boric acid treatments. These were kept in the same growth chambers until sample harvest.

For seedlings grown on plates, sterilized seeds were germinated on 1/4-strength Murashige and Skoog (MS) agar medium (Murashige and Skoog, 1962). 8-day-old seedlings were transferred to 1/4-strength MS medium with different concentrations of boric acid as indicated in Figure 1 and grown in the same growth chamber as described for 4-week old plants.

### Measurement of chlorophyll and root length

Chlorophyll concentrations were determined on a fresh-weight basis. Leaves of 4-week-old *S. parvula* and *A. thaliana* plants were harvested and weighed. Total chlorophyll was extracted with dimethyl sulfoxide solvent (VWR, Radnor, PA), and measured using a SmartSpec™ Plus spectrophotometer (Bio-Rad, Hercules, CA) as described (Richardson et al., 2002). Four biological replicates were used for control and treatments.

Root length was measured daily for 7 days for seedlings grown vertically on 1/4-strength MS agar plates. Root length was measured by marking root tip positions daily at the same time for 7 days for both species. On day 7, the plates were scanned and the root lengths quantified using ImageJ (Schneider et al., 2012). Four biological replicates were used with at least 8 seedlings per replicate.

### Elemental analysis

Shoot and root tissues were harvested at 3 or 24 hours following control (mock) and 5 and 10 mM boric acid treatments. Samples were dried at 37°C for one week in a desiccator to yield between 5 and 60 mg of dry tissue. For quantification of cell wall elements, we prepared cell walls as an alcohol insoluble residue (AIR) as described (Pettolino et al., 2012). Briefly, the harvested tissues were ground to a powder and washed with aq. 80% ethanol, acetone, and methanol. Selected elements (Li, B, Na, Mg, Al, P, S, K, Ca, Fe, Mn, Co, Ni, Cu, Zn, As, Se, Rb, Sr, Mo, and Cd) were quantified using inductively coupled plasma mass spectrometry (ICP-MS) at the United States Department of Agriculture-Agricultural Research Service (USDA-ARS)-Plant Genetics Facility at the Donald Danforth Plant Science Center as described (Baxter *et al*. 2014). Four to five biological replicates were used for each data point. All measurements were normalized to amount per unit weight. One-way ANOVA followed by Tukey’s post-hoc tests implemented in R were used to identify significant differences between samples.

A second independent ICP-MS analysis with a modified protocol that included a rigorous digestion step was conducted to quantify the boron content in the cell walls and to confirm results obtained in the first ICP-MS quantification. AIR from a second set of four biological replicates were prepared as described below. The AIR was digested with 1 mL ultrapure 70% nitric acid (BDH Aristar® Ultra, VWR, Radnor, PA) in 15 mL Teflon beakers, followed by a serial digestion with 100 μL 70% nitric acid on a 100 °C hot plate overnight. Digestions were carefully dried down to almost complete dryness between each step. Acid washed teflon beakers and trace metal clean tubes were used instead of standard laboratory glassware that contain borosilicate in order to minimize the boron background. A final digestion was performed with 100 μL 70% nitric acid and 50 μl 35% H_2_O_2_ (ACS grade, Ward’s Science, Rochester, NY) since undissolved particles remained in solution at the end of the second digestion. The samples were dried on a hot plate as described earlier and the residue dissolved in 5 mL 2% nitric acid. The solution was sonicated for 3-5 mins using an ultrasonic cleaner (FS220, Thermo Fisher Scientific, Waltham, MA). The solution was diluted to 10 mL with 2% nitric acid. Boron was quantified using a Thermo iCap Qc ICP-MS (Thermo Fisher Scientific Inc., Waltham, MA). Internal standard solutions containing ^6^Li, ^45^Sc, ^89^Y, ^103^Rh, ^115^In, ^193^Ir, ^209^Bi were added prior to analysis via a Y-split. Quantification was performed using commercially available standards (IV-ICPMS-71A, Inorganic Ventures, Christiansburg, VA). Pearson correlation coefficient between the two independent ICP-MS experiments was computed using the cor.test function in R.

### Identification of orthologs between *S. parvula* and *A. thaliana*

Genome annotations for *S. parvula* version 2.2 (https://phytozome-next.jgi.doe.gov/) and *A. thaliana* genome version 10 (https://www.araport.org/) were used for ortholog identification. When multiple spliced forms existed in *A. thaliana*, the longest version was considered. Orthologous gene pairs as best reciprocal hits between these two species were identified using the CLfinder-OrthNet pipeline with default settings (Oh and Dassanayake, 2019). To account for lineage-specific gene duplications in both species, orthologous gene pairs were searched reciprocally between the two-species using BlastP with an e-value of 1e-5 and MMseqs2 (Steinegger and Söding, 2017) with an equivalent e-value cutoff. These pairs were further filtered using OrthoFinder (Emms and Kelly, 2015) with granularity -I of 1.6 and were added back to the CLfinder pipeline to extract all possible ortholog pairs between the two species. Among a total of 27,206 *A. thaliana* protein-coding gene models, 22,112 were paired with at least one *S. parvula* homolog. Similarly, 21,673 out of 26,847 *S. parvula* gene models were paired with at least one *A. thaliana* ortholog. The two reciprocal searches were merged, and redundant pairs were removed to generate 23,281 *S. parvula*-*A. thaliana* orthologous gene pairs.

### Transcriptome profiling

Root and shoot tissues were harvested separately for each plant 24 hours after boric acid treatment. Total RNA (at least 6 μg) was extracted using the RNeasy Plant Mini kit (Qiagen, Hilden, Germany), with an additional step to remove contaminating DNA. Four biological replicates per condition were generated and three were used for RNA-seq libraries. RNA-seq libraries were prepared with a TruSeq Stranded mRNAseq Sample Prep kit (Illumina, San Diego, CA, USA) at the Roy J. Carver Biotechnology Center, University of Illinois at Urbana-Champaign. Libraries were barcoded and sequenced on three lanes of HiSeq2500 platform (Illumina), generating > 25 million high-quality 100-nucleotide (nt) single-end RNA-seq reads per sample. These reads are deposited in the BioProject PRJNA663969 at the NCBI-SRA database.

RNA-seq reads after quality checks using FastQC (https://www.bioinformatics.babraham.ac.uk/projects/fastqc/) from each sample were mapped to either *A. thaliana* TAIR10 or *S. parvula* genome v2 using HISAT2 version2.0.1(Kim et al., 2015) with default parameters. A custom Python script was used to count uniquely mapped reads to each gene model found to be expressed. To identify a list of robust differentially expressed genes (DEGs), we used a consensus list from DEGs identified using a parametric method, DESeq2 (Love et al., 2014) and a non-parametric method, NOISeq (Tarazona et al., 2015) with a FDR-adjusted p-value cutoff set to 0.05. Only genes selected by both methods as significantly different were used for down-stream analyses.

To compare the expression levels of orthologs between *S. parvula* and *A. thaliana*, expression values of reads per kilobase of transcript per million mapped reads (RPKM) < 1 were removed. The RPKM values of filtered ortholog pairs were converted to log_2_-transformed counts and median-normalized. These normalized RPKM values for ortholog pairs across all samples from shoots and roots were subjected to fuzzy k-means clustering (Gasch and Eisen, 2002) to identify co-expressed gene groups. Ortholog pairs in each of the resulting clusters were further filtered based on: (1) the membership of a given ortholog pair was no less than 0.5; (2) the expression changes of each ortholog pair in a given cluster were considered to be statistically significant by both DESeq2 and NOISeq to ensure that the expression pattern of a given pair agreed with that for the cluster; and (3) clusters in which the pattern was consistent between all biological replicates were considered for downstream analyses.

BiNGO (Maere et al., 2005) was used to identify enriched networks of Gene Ontology (GO) terms in each species. To reduce the redundancy between enriched GO terms and their associated inference related to DEGs, redundant GO terms with > 50% overlap with similar terms were further clustered using Markov clustering implemented via GOMCL (https://github.com/Guannan-Wang/GOMCL) (Wang et al., 2020). Custom Python scripts were used to extract all direct child terms of a given GO term or all genes annotated with the given GO term. GO terms with zero assigned genes from *A. thaliana* were removed from the analysis. Kyoto Encyclopedia of Genes and Genomes (KEGG) (Kanehisa et al., 2016) was used to map genes to specific metabolic pathways.

### RT-qPCR

Plants were grown and treated with excess boric acid and harvested as described for RNA-seq experiments. Total RNA (0.5 μg) was used in a 20 μL reverse transcription (RT) reaction for first-strand cDNA synthesis with SuperScript™ III Reverse Transcriptase according to the manufacturer’s instructions (Invitrogen, Carlsbad, CA). The reverse transcription products were diluted to 200 μL, and 2 μL was used in a 20 μL qPCR reaction using SYBR™ Select Master Mix (Applied Biosystems, Foster City, CA) in a ViiA 7 Real-Time PCR System (Applied Biosystems, Foster City, CA).

Our transcriptomic profiles showed that the commonly used reference genes, *ACT2*, *CYTC-1*, *CYTC-2*, *EF1α*, *UBQ10*, and *GAPDH*, all had expression levels that varied between treatments, plants, and tissue types, which made them unsuitable as reference genes in RT-qPCR. Therefore, we searched for the most uniformly expressed and conserved genes in both plants and selected At5g46630 (ADAPTOR PROTEIN-2 MU-ADAPTIN) and At4g26410 (RGS1-HXK1 INTERACTING PROTEIN 1) and their *S. parvula* orthologs as internal reference genes, following the best practice recommendations from previous studies (Czechowski et al., 2005; Wang et al., 2014). RT-qPCR primer sequences are listed in Supplemental Data Set 5.

### Metabolomics analyses

Shoot and root samples were harvested at 24 hours after control, 5, and 10 mM boric acid treatments and freeze dried (FreeZone 2.5 Plus, Labconco Corp., Kansas City, MS). Untargeted profiling of polar metabolites, including boric acid, using gas chromatography-mass spectrometry (GC-MS) was performed at the Metabolomics Center at University of Missouri, Columbia. To facilitate the detection of trace metabolites, 20 mg of roots or 50 mg of shoots from pools of 7-12 plants were used per biological replicate per condition. The dry tissues were suspended in 1.0 ml of aq. 80% methanol and 20 μl of HPLC grade water containing 1 μg/ml ribitol. The suspensions were vortexed for 20 seconds, and sonicated for 15 min. The suspensions were shaken for 2 hours at 140 rpm in an orbital shaker and centrifuged for 30 minutes at 15000 g. Equal amounts of the supernatant were transferred to autosampler vials. The solutions were concentrated to dryness using a gaseous nitrogen stream. The dried extracts were methoximated with 25 μl of 15 mg/mL methoxyamine hydrochloride in pyridine, and trimethylsilylated with 25μL N-methyl-N-(trimethyl-silyl)trifluoroacetamide (MSTFA) and 1% chlorotrimethylsilane (TMCS). The derivatized extracts were analyzed for non-targeted metabolic profiling using an Agilent 6890 GC coupled to a 5973N MSD mass spectrometer with a scan range from m/z 50 to 650 (Agilent Technologies, Inc., Santa Clara, CA). 1 μl of sample was injected into the GC column with a split ratio of 1:1 for polar GC-MS analysis. Separation was achieved using a 60 m DB-5MS column (J&W Scientific, 0.25 mm ID, 0.25 um film thickness) with a temperature program of 80 °C for 2 min, then ramped at 5 °C /min to 315 °C and held at 315 °C for 12 min, and a constant flow of helium gas (1.0 ml/min). A standard alkane mix was used for GC-MS quality control and retention index calculations. The raw data were first deconvoluted using AMDIS software (http://chemdata.nist.gov/dokuwiki/doku.php?id=chemdata:amdis) and annotated through mass spectral and retention index matching to an in-house spectra library. The unidentified compounds were searched and identified using spectral matching to a commercial NIST17 mass spectral library. The raw abundance/intensity for each identified compound was normalized with the internal standard, ribitol (peak area of each metabolite/peak area of internal standard × 1,000). Different molecular features were manually curated to the most relevant molecular feature for each identified metabolite. A minimum of three biological replicates were used for each condition for each tissue. Significant differences between samples were determined by Student’s t tests followed by FDR correction for multiple testing (Benjamini and Hochberg, 1995) in MetaboAnalystR (Chong et al., 2019).

### Yeast complementation assay

Yeast *Δbor* mutant strains were in the *MATα ADE2 his3*Δ*1 leu*Δ*D0 lys2*Δ*0 TRP1 ura3*Δ*0 bor1D::KanMX* background. The entire coding regions of *AtBOR4*, *AtBOR5*, *SpBOR4*, *SpBOR5* were cloned from *A. thaliana* and *S. parvula* cDNA, respectively, and were separately introduced into the pDD506 plasmid (Wang and Donze, 2016), driven by the ADH1 promoter. The *Δbor* mutant was transformed with the recombinant pDD506 plasmids or the empty pDD506 plasmid as a negative control. The transformants were selected on SD medium-His. Boron toxicity tolerance assays were performed as described in Nozawa *et al*. (2006). Briefly, yeast cells were grown in SD medium to OD_600_=1, collected, and spotted onto solid SD or SD containing 80 mM boric acid with different titers. The plates were photographed after incubation for 10 days at 30 °C.

## Supplemental Data

**Supplemental Figure 1.**
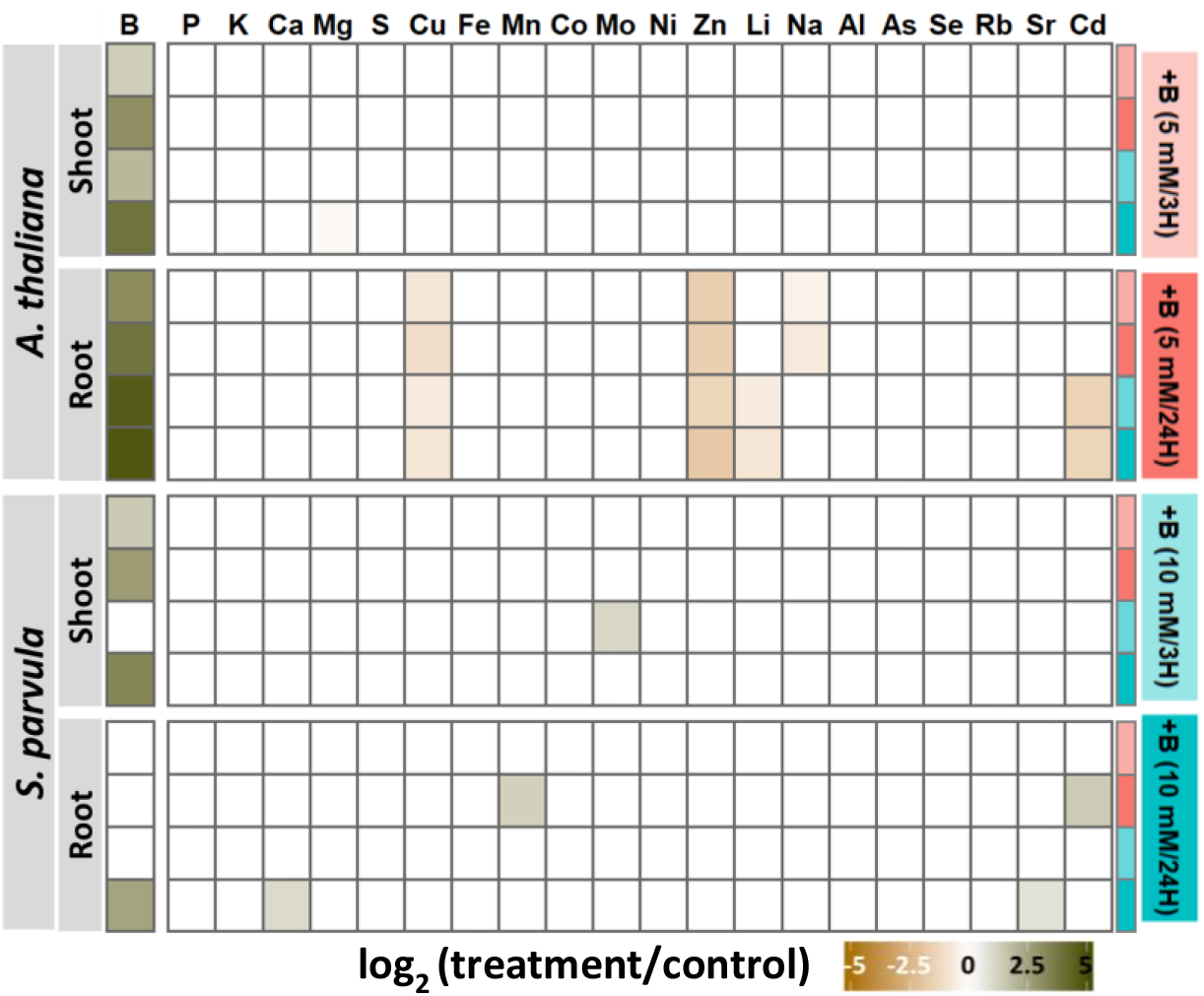
Ionomic profiles in response to boric acid treatments. Significant differences of each treatment compared to control were based on one-way ANOVA followed by Tukey’s post-hoc tests (p < 0.05).

**Supplemental Figure 2.**
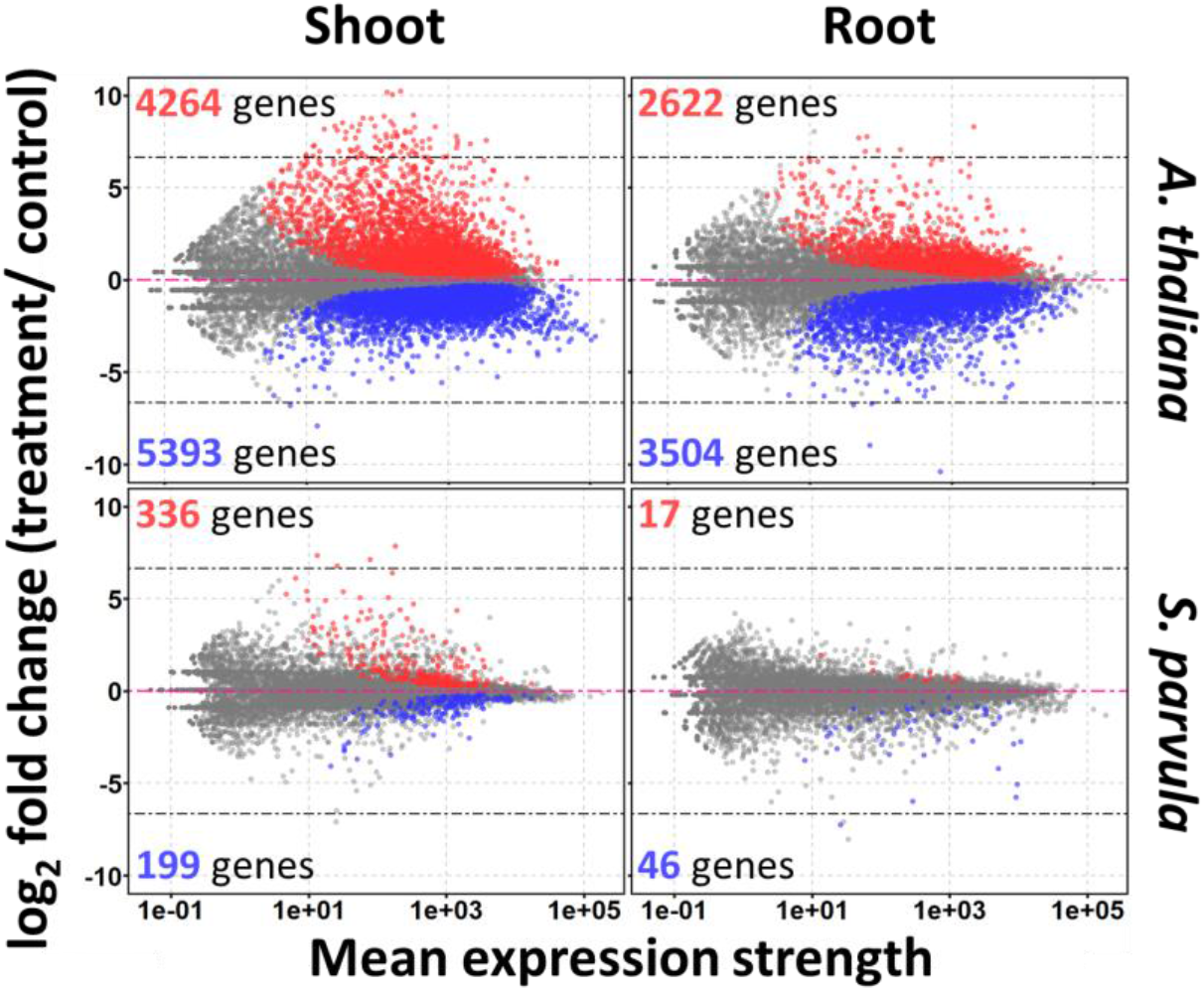
Differential expression visualized using MA-plots from shoot (left panel) and root (right panel) of *A. thaliana* (upper panel) and *S. parvula* (lower panel). Red dots represent up-regulated DEGs and blue dots indicate down-regulated DEGs.

**Supplemental Figure 3.**
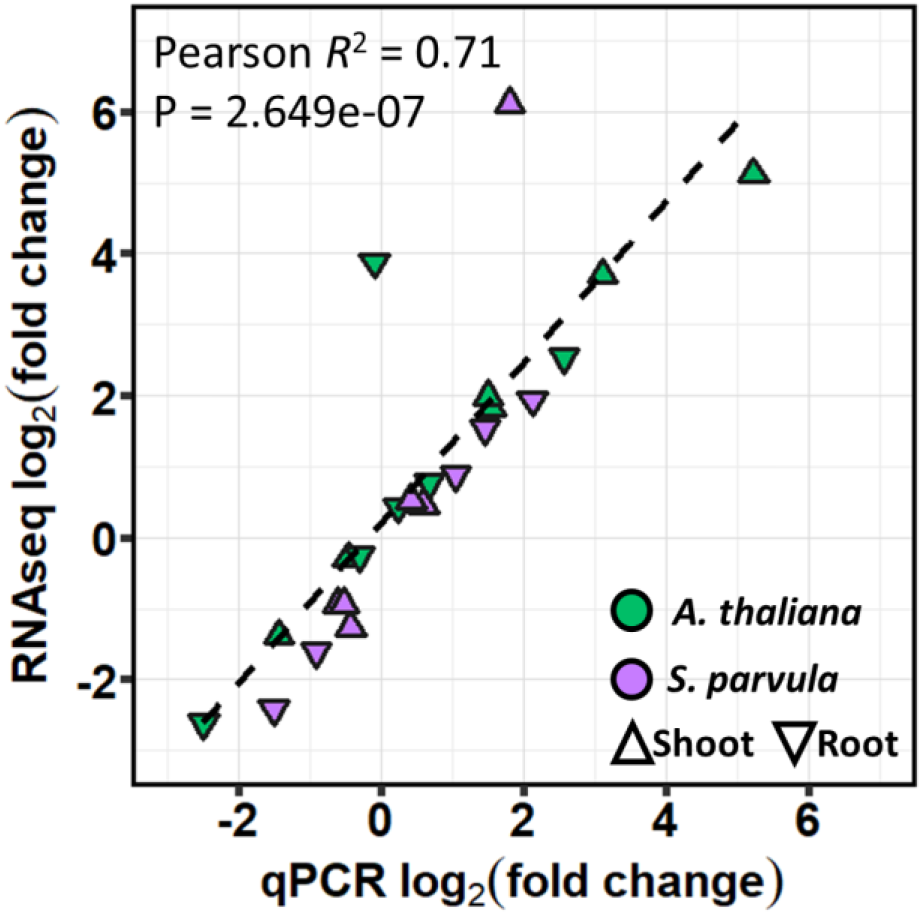
Assessment of qPCR and RNA-seq expression data agreement for selected differentially expressed genes.

**Supplemental Figure 4.**
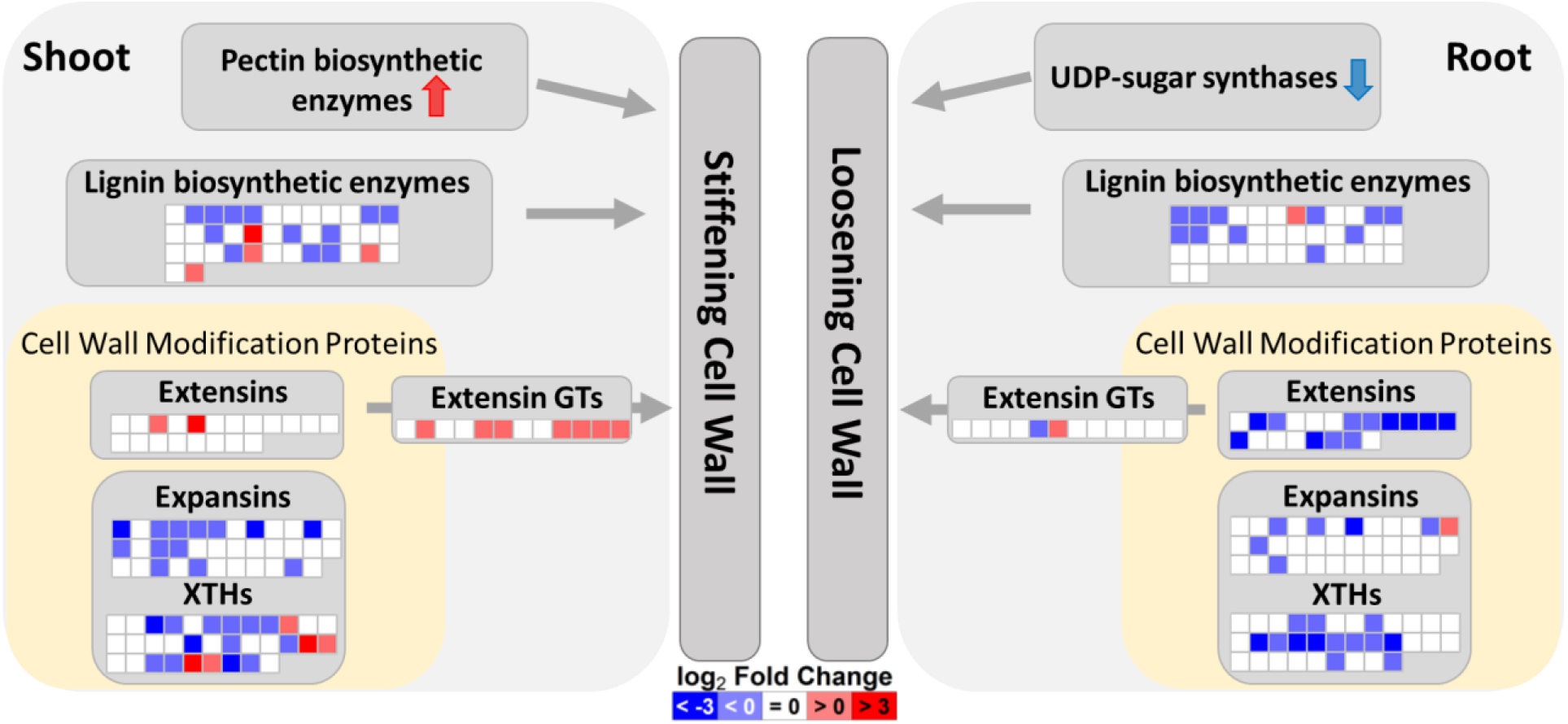
Summary of cell wall modifications in *A. thaliana* in response to boric acid treatment. Genes are represented by in colored blocks, grouped into families and pathways, with up- and down-regulation marked by red and blue, respectively.

**Supplemental Figure 5.**
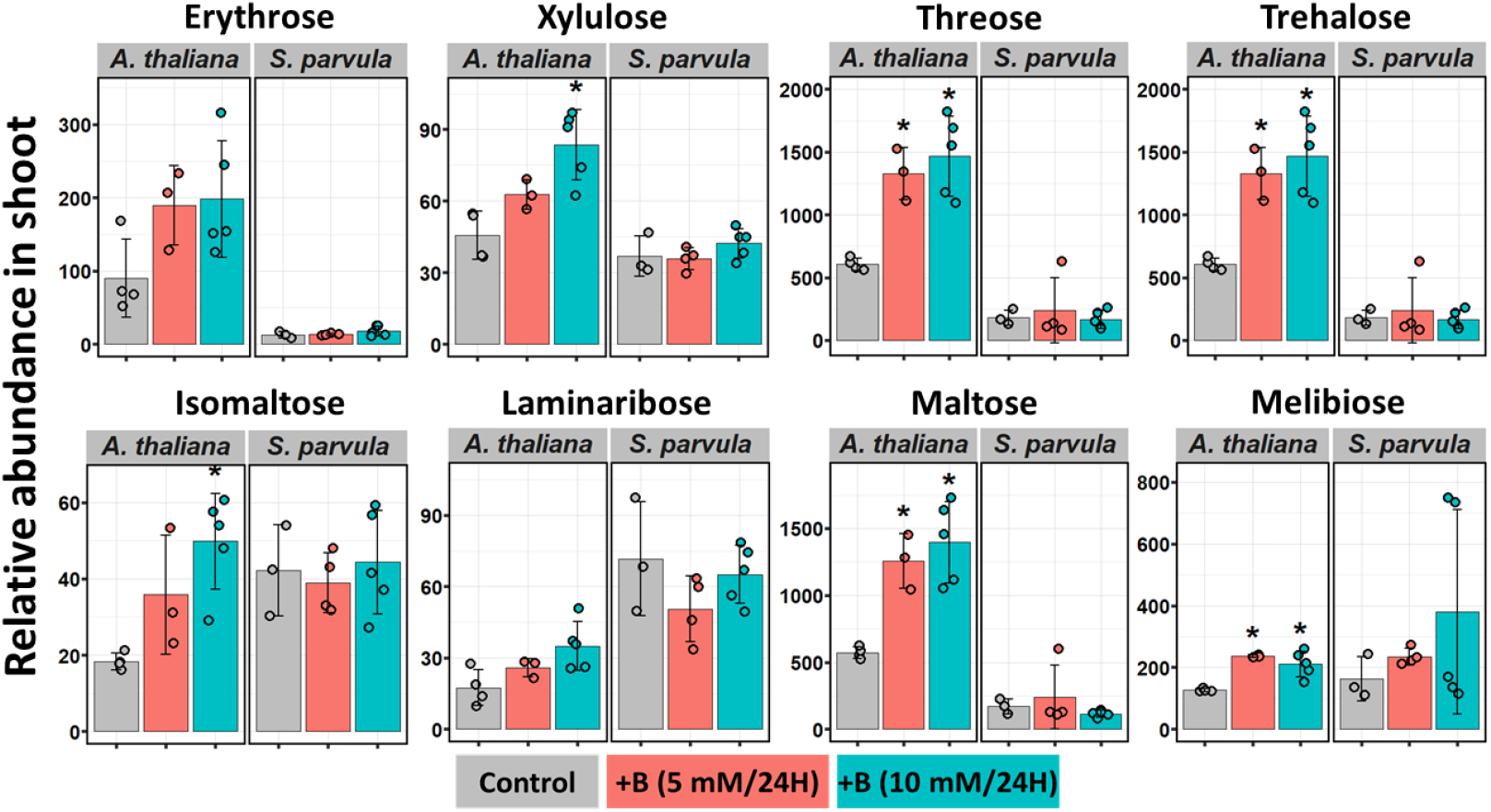
Relative abundances of sugars in *A. thaliana* shoots that are not directly associated with cell wall polysaccharides. The relative abundance is given compared to the internal standard, ribitol. Values shown are mean ± SD (n = 3, 4 or 5). Asterisks represent significant differences of each treatment compared to control according to Student’s t test (p < 0.05).

**Supplemental Figure 6.**
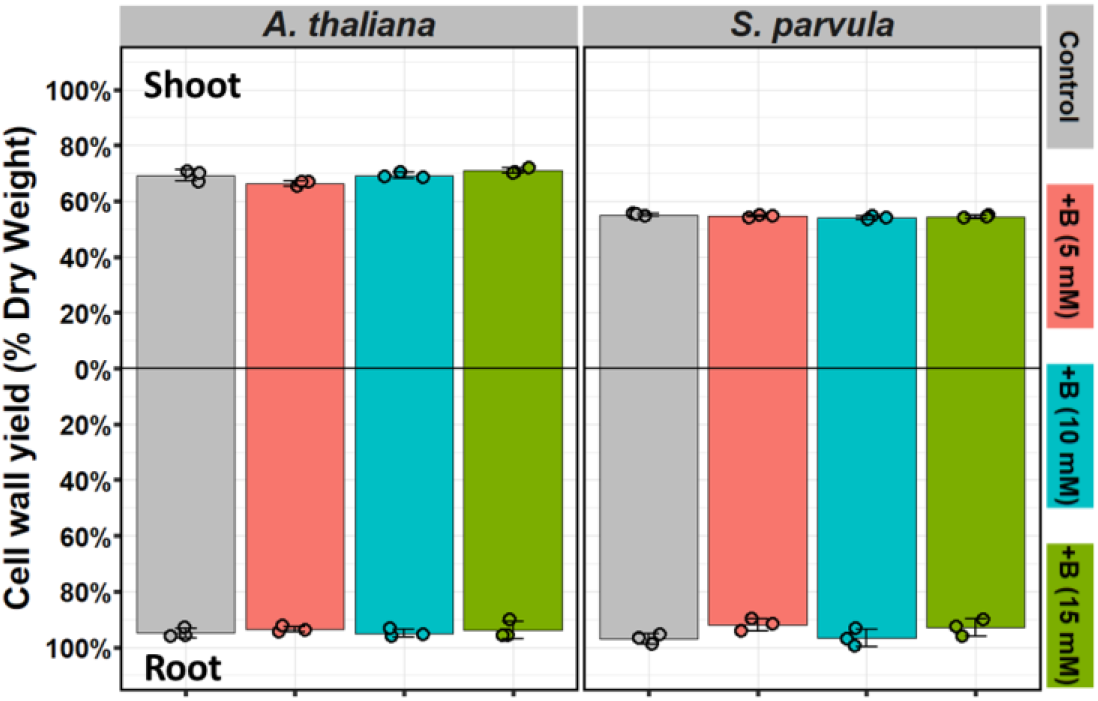
Cell wall content in *A. thaliana* and *S. parvula* treated with excess boron for 5 days. Values shown are mean ± SD (n = 3 or 4).

**Supplemental Figure 7.**
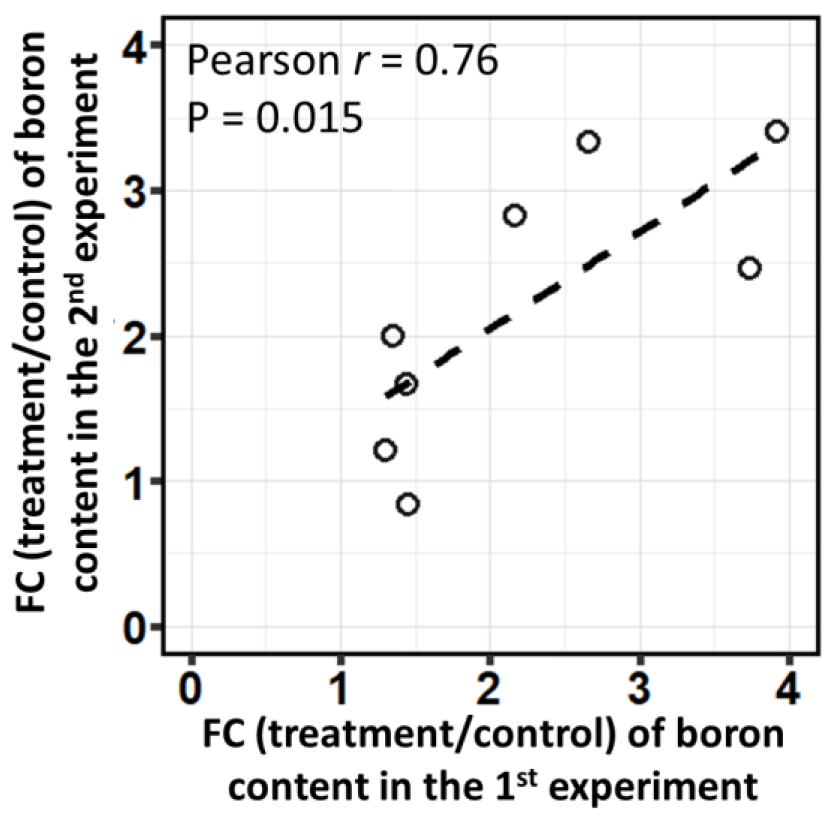
Conformance of boron content quantified using two independent experiments. Fold changes were calculated comparing the treatment to the control for each tissue from each species. Pearson’s *r* and *p*-values are indicated.

**Supplemental Figure 8.**
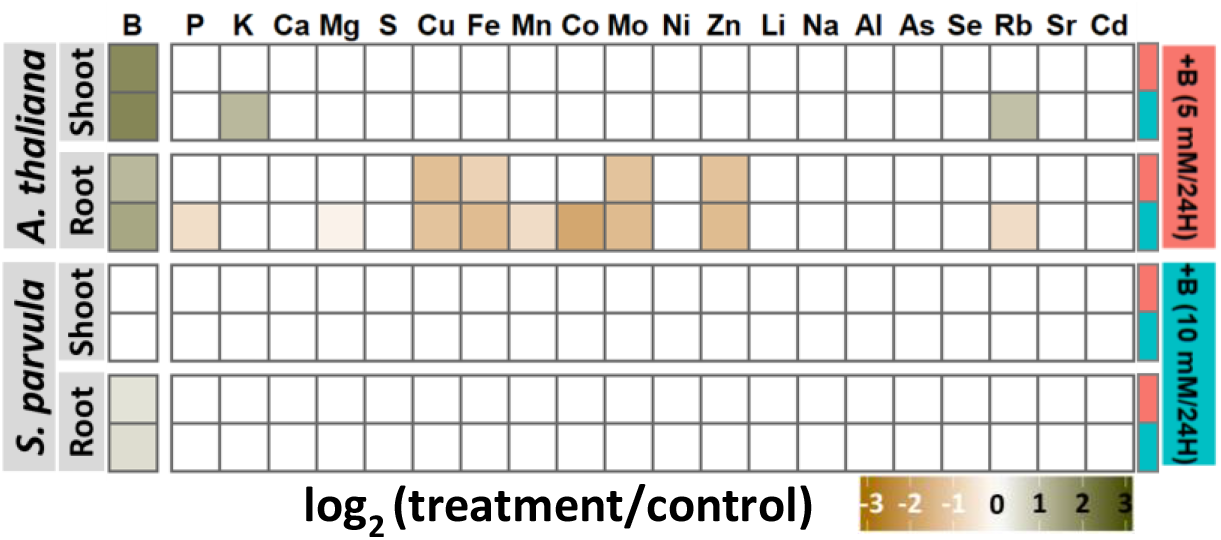
Ionomic profiles in the cell wall extracts in response to boric acid treatments at 24 hours. Significant differences of each treatment compared to control for each element were based on one-way ANOVA followed by Tukey’s post-hoc tests (p < 0.05).

**Supplemental Figure 9.**
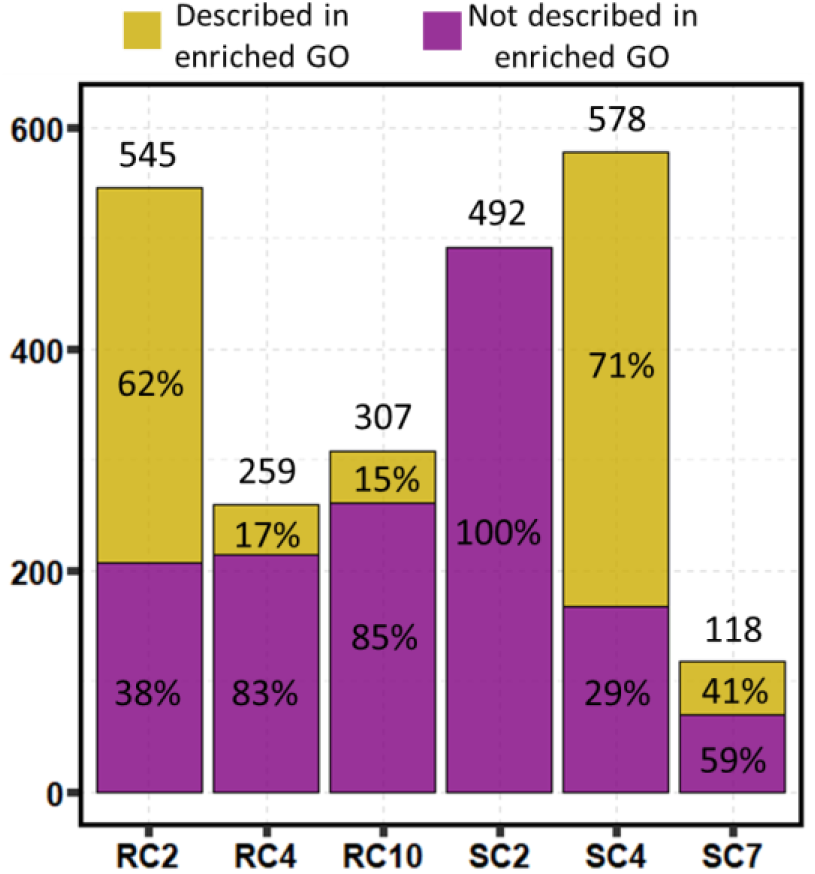
Number of members and their annotation availability in selected stress-ready clusters. RC: root cluster; SC: shoot cluster.

**Supplemental Figure 10.**
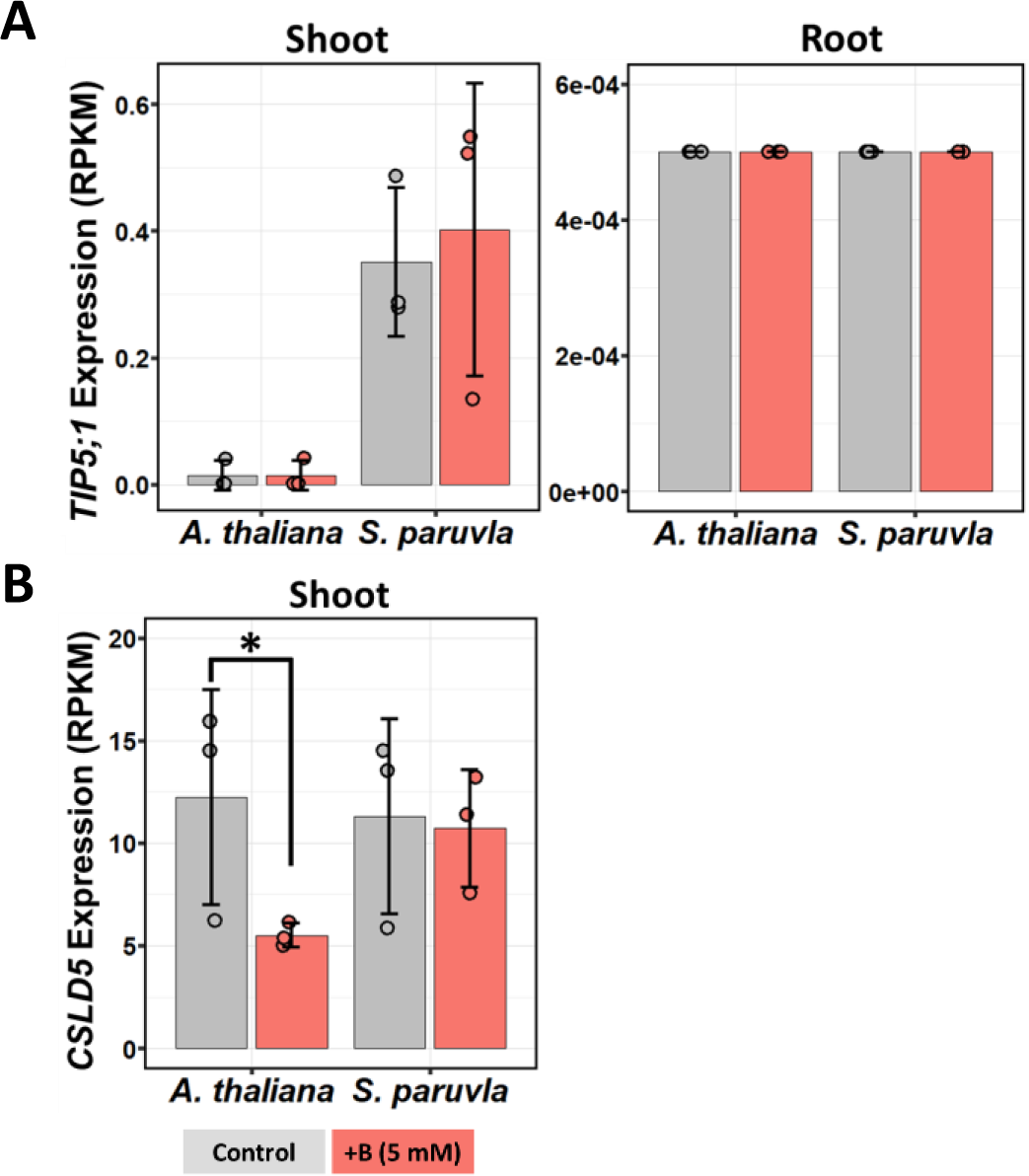
Expression levels of *TIP5;1* and *CSLD5* from *A. thaliana* and *S. parvula* in control and 5 mM boric acid treatment. Asterisks represent significant differences in expression compared to control (at FDR-adjusted p<0.05) determined by both DESeq2 and NOISeq.

**Supplemental Figure 11.**
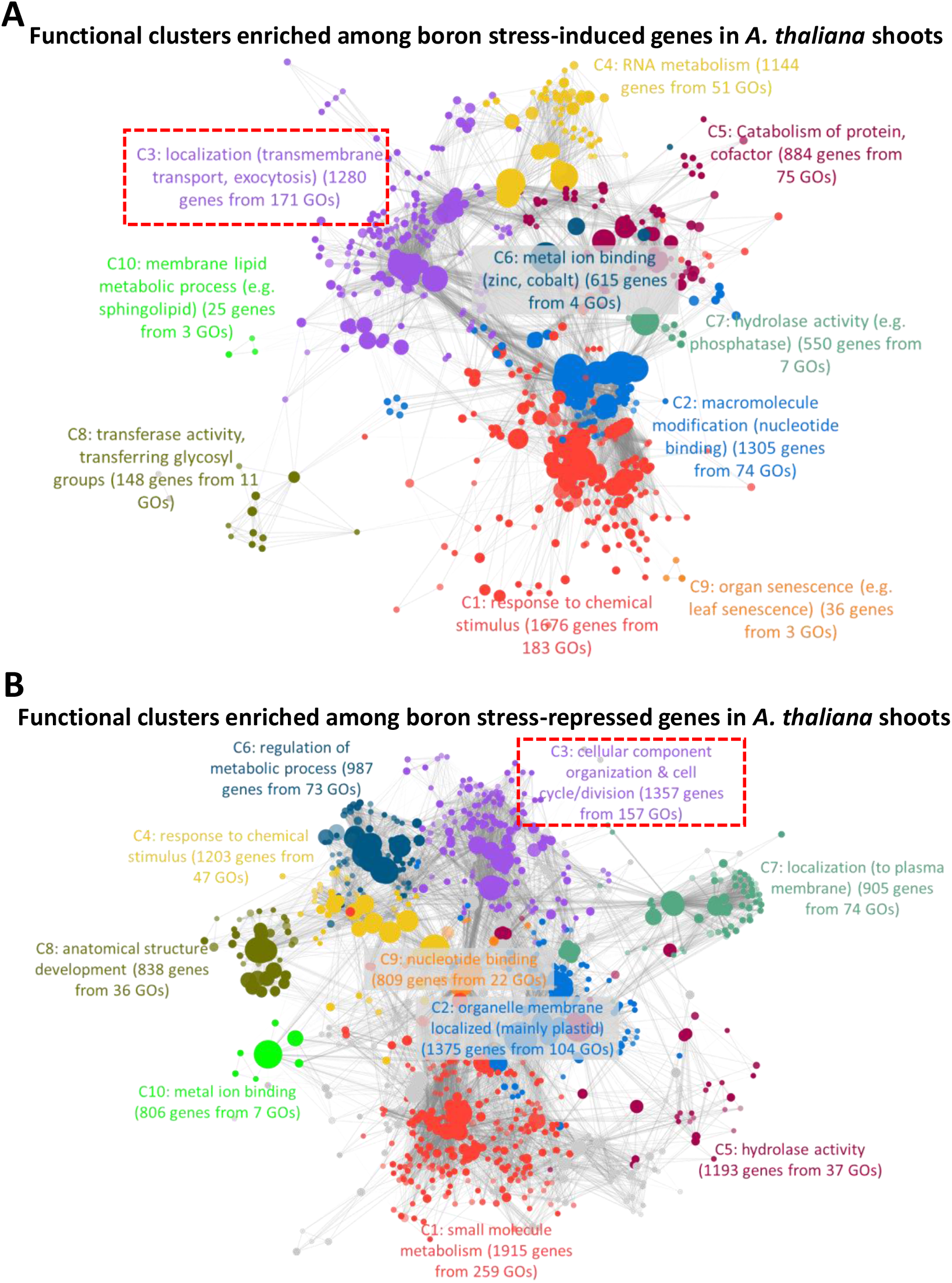
Functional clusters enriched among genes that were induced (A) and repressed (B) in *A. thaliana* shoots. Top 10 largest clusters are differently colored and labelled with the representative functional terms. Each node represents a GO term; node size represents genes in the test set assigned to that functional term; GO terms sharing more than 50% of genes are connected with edges; and shade of each node represents the *p*-value assigned by the enrichment test (FDR-adjusted p<0.05) with darker shades indicating smaller *p*-values.

**Supplemental Data Set 1**. The list of DEGs identified from DESeq2 and NOIseq, and list of genes used for pathway analysis.

**Supplemental Data Set 2**. Functional clusters enriched in boron stress-responsive genes in *A. thaliana* and *S. parvula*.

**Supplemental Data Set 3**. List of differentially accumulated metabolites and their functional categories.

**Supplemental Data Set 4**. Co-expression clusters of orthologs between *A. thaliana* and *S. parvula*.

**Supplemental Data Set 5**. Primer sequences used for qPCR.

## Acknowledgements

This work was supported by the National Science Foundation awards MCB-1616827 and NSF-IOS-EDGE-1923589 to MD, DHO, and APS. MD also acknowledges the support from the Next-Generation BioGreen21 Program of Republic of Korea (PJ01317301). GW was supported by an Economic Development Assistantship award from Louisiana State University. MAO acknowledges the Division of Chemical Sciences, Geosciences, and Biosciences, Office of Basic Energy Sciences of the United States Department of Energy through Grant DE-SC0008472 for funding studies of plant cell walls. DMC acknowledges NSF through award MCB-1818312. The authors thank Dr. David Donze (Biological Sciences, LSU) for providing necessary yeast strains and advice on the yeast assays and Katherine Winchester (high school student, St. Joseph’s Academy, Baton Rouge) for her assistance with root phenotyping. The authors also acknowledge the LSU High Performance Computing services for providing computational resources needed for data analyses.

## Author contributions

GW and MD developed the experimental design; GW prepared plant samples, conducted data analyses, and performed RT-qPCR; SFD performed yeast assays; ADH and GW designed and conducted the ICP-MS assays. GW, DHO, DMC, MAO, APS, and MD contributed to data interpretation. GW and MD wrote the article with input from all co-authors who revised and approved the final manuscript.

## Additional information

Correspondence and requests for materials should be addressed to MD.

